# In Situ Photoactivated Hydrogel Adhesive Dressings for Post Colon Polypectomies (PolypCures)

**DOI:** 10.64898/2025.12.11.693663

**Authors:** Xingyu Hu, Carolina Villarreal-Otalvaro, Rong Liu, Yin P. Hung, Shawna Pratt, George A. O’Toole, Samantha Berry, Michal Jones, Aaron Rosenthal, Mark W. Grinstaff

## Abstract

Colon polypectomy is a widely performed endoscopic procedure that reduces the incidence of colorectal cancer but leaves exposed colonic wounds susceptible to bleeding, perforation, and infection. The current standard of wound care, mechanical clips, is limited by technical complexity, high cost, and poor efficacy for large (> 2 cm in diameter) or difficult-to-access lesions. Here, we present PolypCure, an *in situ* photoactivated hydrogel adhesive dressing delivered via a single catheter, through either oozing or spraying methods, to achieve rapid, spatiotemporally controlled sealing and hemostasis (within 2 min) of colon lesions using the white light source integrated into standard endoscopic instruments. The hydrogel, composed of norbornene- and thiol-functionalized polyethylene glycol, carboxymethyl cellulose, and Eosin Y, exhibits mechanical properties comparable to colon tissue (G’ ∼ 6 kPa), strong shear strength (> 15 kPa), low swelling (< 200%), and small pore size (∼ 20 µm), ensuring long-term stability (up to 30 days) and protection against bacterial infiltration. The formulation also demonstrates biocompatibility and hemocompatibility. In an *in vivo* pig colon endoscopic mucosal resection (EMR) model, PolypCure fully adheres to large lesions for at least 3 days and promotes early-stage wound healing. Overall, PolypCure is a promising solution to wound management post-polypectomy.

## 1. Introduction

Colorectal cancer remains a leading cause of mortality and morbidity worldwide, with a 5-year survival rate of approximately 65%.^[1–3]^ It typically arises from polyps – abnormal growths on the inner lining of the colon or rectum. Advances in early detection and therapeutic interventions, particularly colonoscopic polypectomy, significantly reduce colorectal cancer incidence and mortality.^[4,5]^ In the United States alone, over 15 million colonoscopies are performed annually, with precancerous polyps identified in 25% of cases, often requiring removal.^[6]^

Colon polypectomy involves endoscopic mucosal resection (EMR) or endoscopic submucosal dissection (ESD) to remove early-stage cancerous tumors or benign lesions from the gastrointestinal (GI) tract.^[7–10]^ For lesions ≤ 2 cm in diameter, EMR is the standard, whereas ESD enables en-bloc removal of larger lesions.^[8,10]^ Although these techniques are minimally invasive and generally safe,^[11,12]^ they carry risks for adverse events, including intraprocedural bleeding, delayed bleeding, delayed perforation, and, in severe cases, sepsis.^[13–15]^ Intraprocedural bleeding occurs in 1.5% to 2.8% of polypectomies and can persist for up to 2 days.^[16,17]^ Delayed bleeding, the most frequent complication, typically occurs 2-7 days post-procedure, affecting 1-7% of EMR and up to 15% of ESD cases.^[16–20]^ Risk factors such as lesion size, hepatic flexure proximity, and anticoagulant use further increase this rate to 17.2%.^[21–23]^ Less commonly, but more severely, delayed perforation occurs in up to 5% of colonoscopic interventions and usually arises within 24 hours.^[24,25]^ While intraprocedural events are often managed endoscopically, delayed bleeding and perforation are more challenging to detect and treat promptly, potentially resulting in severe blood loss, emergency surgery, or additional hospitalizations.^[7,10,26]^

The current standard of care for managing post-polypectomy wounds is prophylactic clipping. While clips are effective for acute hemostasis,^[27]^ their ability to prevent delayed bleeding and perforation is inconsistent and highly dependent on lesion type, location, and operator skill.^[11,17,28–30]^ Moreover, clip placement is technically demanding, time-consuming, costly, and may cause secondary mucosal injury.^[15,26,31]^ Several products developed for other indications, such as hemostatic sponges and gels (e.g., Surgicel, Gelfoam, Fibrillar), hemostatic powders (e.g., Hemospray, Endoclot, Ankaferd Blood Stopper), or fibrin sealants (Tisseel) are off-label options to treat post-polypectomy wounds.^[7,17,32]^ However, their effectiveness remains limited due to weak mucosal adhesion, rapid degradation, and/or the potential for infection.^[33]^ Hemostatic powders, in particular, are difficult to apply endoscopically and exhibit short residence times (< 48 hr), with reported rebleeding rates of 11-23%.^[17]^ PuraStat (3D Matrix Ltd., France), a self-assembling peptide gel, is currently the only topical hemostat approved specifically for post-EMR/ESD use.^[31]^ While it forms a transparent gel upon contact with biological fluids and reduces intraprocedural and delayed bleeding in some studies,^[34–36]^ its weak mechanical strength and poor adhesion limit its performance, especially on inflamed or irregular tissue surfaces.^[17,37]^ Currently, no clinically approved solution offers strong mucosal adhesion, broad applicability across wound sizes, effective support for wound healing, and reliable prevention of delayed complications following colon polypectomy.

Here, we introduce PolypCure, an *in situ* photoactivated hydrogel dressing delivered through a standard endoscopic catheter immediately following polypectomy (Fig 1A). The hydrogel is formed through visible-light–induced thiol–norbornene step-growth photopolymerization between norbornene (NB)- and thiol (SH)-functionalized four-arm polyethylene glycol (PEG) macromers,^[38]^ with carboxymethyl cellulose (CMC) incorporated as a thickener and secondary adhesive component (Fig 1B). Upon illumination with the white light integrated into standard endoscopes, the precursor solution rapidly crosslinks within 2 minutes to form a cohesive, adherent barrier that seals submucosal wounds (Fig 1C). In a large animal porcine model, PolypCure achieves complete coverage and adhesion on large colon lesions (> 2 cm) for at least 3 days without adverse effects, reducing inflammation and necrosis while promoting fibroplasia and re-epithelialization. These results establish PolypCure as a practical and effective platform for post-polypectomy wound management, with strong potential to reduce complications and improve clinical outcomes.

**Figure 1.**
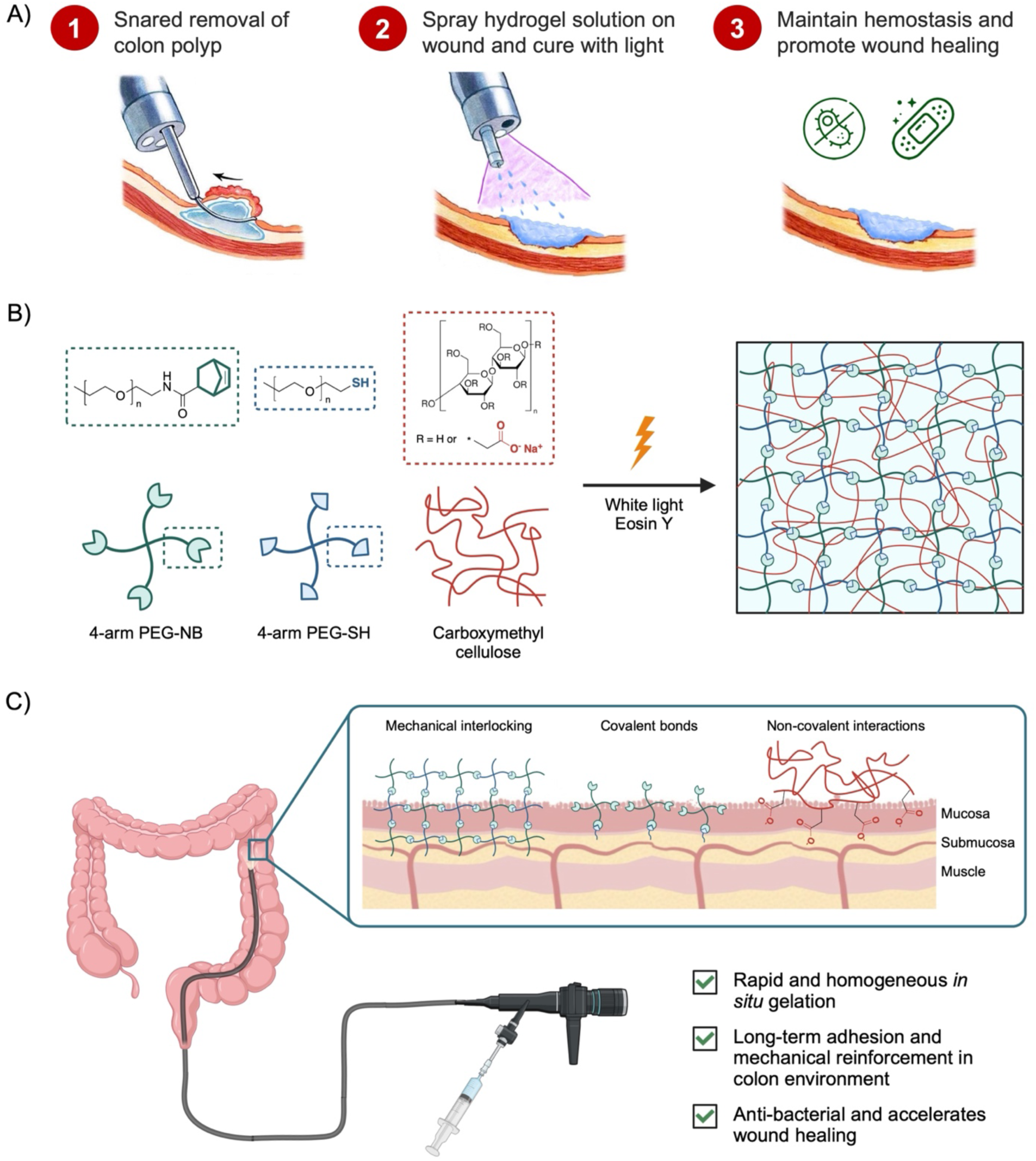
Schematic illustration of the PolypCure hydrogel adhesive dressing, describing its composition, crosslinking mechanism, tissue adhesion process, and role in wound protection post colon polypectomy. A) Following colon polyp resection, the hydrogel precursor solution is delivered through a standard catheter to coat the wound site and is immediately cured using the endoscope’s built-in white light, ensuring hemostasis and promoting wound healing. B) Upon white light irradiation, 4-arm PEG-norbornene (NB), 4-arm PEG-thiol (SH), and carboxymethyl cellulose (CMC) undergo crosslinking to form a gel network. C) The hydrogel adheres to the colon mucosa and submucosa through: 1) mechanical interlocking, facilitated by polymer diffusion during in situ gelation, 2) covalent bonding between 4-arm PEG-NB and tissue thiols, and 3) non-covalent interactions, including hydrogen bonding and physical entanglements.

## 2. Results and Discussion

### 2.1. Design and characterization of the PolypCure hydrogel

We developed the PolypCure system to meet seven functional design criteria essential for post-polypectomy wound management: 1) easy application using standard clinical instrumentation; 2) precisely targeting and complete coverage of lesions of all sizes and locations; 3) rapid gelation, wound closure, and hemostasis (< 5 min); 4) strong adhesion and mechanical properties comparable to native colon tissue; 5) sustained adhesion and stability in the colonic environment (> 7 days); 6) homogeneous hydrogel formation over the wound to prevent bacteria contamination; and, 7) biocompatibility and hemocompatibility.

To achieve these goals, PolypCure integrates a pre-mixed hydrogel precursor solution with a standard clinical endoscope that serves as both the light source and delivery platform. The precursor formulation consists of 4-arm polyethylene glycol norbornene (PEG-NB), 4-arm polyethylene glycol thiol (PEG-SH), carboxymethyl cellulose (CMC), and the visible-light photoinitiator Eosin Y, with a trace of blue dye for visualization. Delivered through a single-lumen black catheter, the precursor is applied directly to the lesion and rapidly crosslinks under endoscopic illumination, forming a cohesive, adherent hydrogel layer that seals and protects the wound surface.

We selected this step-growth thiol–norbornene chemistry for its compatibility with endoscopic light sources, rapid and controllable in situ gelation, and excellent biocompatibility. This mechanism allows the liquid precursor to conform to irregular wound geometries and partially infiltrate the submucosal surface prior to curing, promoting mechanical interlocking and physical entanglement with the tissue.^[7]^ After gelation, the exposed surface is non-adhesive due to the non-fouling properties of PEG,^[39]^ preventing unwanted bowel adhesion. The orthogonal reaction pathway is expected to produce a highly ideal, homogeneous, densely crosslinked network with small pore size and tunable elasticity—attributes critical for robust adhesion and mechanical stability.^[40,41]^ Unlike free-radical chain-growth polymerization, thiol–norbornene photopolymerization proceeds efficiently in oxygenated and hydrated environments at low photoinitiator concentrations, supporting biocompatibility and hemocompatibility.^[38,41,42]^ Eosin Y is the sole photoinitiator because its strong absorbance in the 400–700 nm range matches the white-light output of standard endoscopes, eliminating the need for ultraviolet activation or cytotoxic co-initiators.^[38,43]^ Finally, we incorporated CMC to increase viscosity and localize the precursor at the wound site during delivery, while also providing secondary non-covalent interactions that enhance toughness and interfacial adhesion.^[44]^

Notably, the PolypCure system employs a standard single-lumen catheter for hydrogel delivery, in contrast to the dual- or tri-lumen systems required by other platforms,^[13,26,27,45]^ significantly simplifying the application procedure. This design enables complete mixing of all components prior to injection, as opposed to *in situ* mixing at the tissue site,^[26,45–47]^ yielding a more homogeneously crosslinked hydrogel with smaller pores, enhanced adhesive strength and improved barrier properties. Furthermore, visible-light activation provides precise spatiotemporal control over gelation, enabling on-demand curing, and facilitating seamless integration into standard endoscopic procedures.

To optimize the formulation, we systematically varied PEG polymer architecture and linkage chemistry, photoinitiator concentration, thickener type and concentration, and dye incorporation for visibility. We also characterized the spectral output and intensity of various endoscope light sources to confirm efficient photoactivation of the hydrogel. We synthesized the 4-arm PEG-NB by reacting excess 5-norbornene-2-carboxylic acid with 5kDa 4-arm PEG-amine through a standard N,N’-dicyclohexylcarbodiimide coupling reaction,^[48,49]^ yielding 95% conjugation as confirmed by NMR spectroscopy (Fig S1, S2). We used the 2-arm ester-linked PEG-SH and 4-arm methylene-linked PEG-SH, functionalized using mercaptopropionic acid,^[50]^ to form the 2-arm and the 4-arm crosslinking systems, respectively (Fig S1, S3).

We first investigated the effect of Eosin Y concentration on gelation time and mechanical properties using the 2-arm system at 20 wt% and a high-intensity fiber-optic light source (AmScope Max). Eosin Y, upon visible light excitation, generates thiyl radicals by abstracting hydrogen from thiol groups, and initiates thiol-norbornene photopolymerization.^[38,43]^ We maintained a 1:1 molar ratio of norbornene to thiol in all formulations to maximize crosslinking efficiency.^[51]^ Escalating Eosin Y concentration from 0.005 mM to 1 mM accelerates gelation and enhances storage modulus (G’) as demonstrated by *in situ* photorheometry, gelation time assays, and rheological measurements (Fig 2A, Fig S4), consistent with trends reported in the literature.^[52]^ At 1 mM Eosin Y, gels form within 10 s and reached a storage modulus (G’) exceeding 20 kPa.

**Figure 2.**
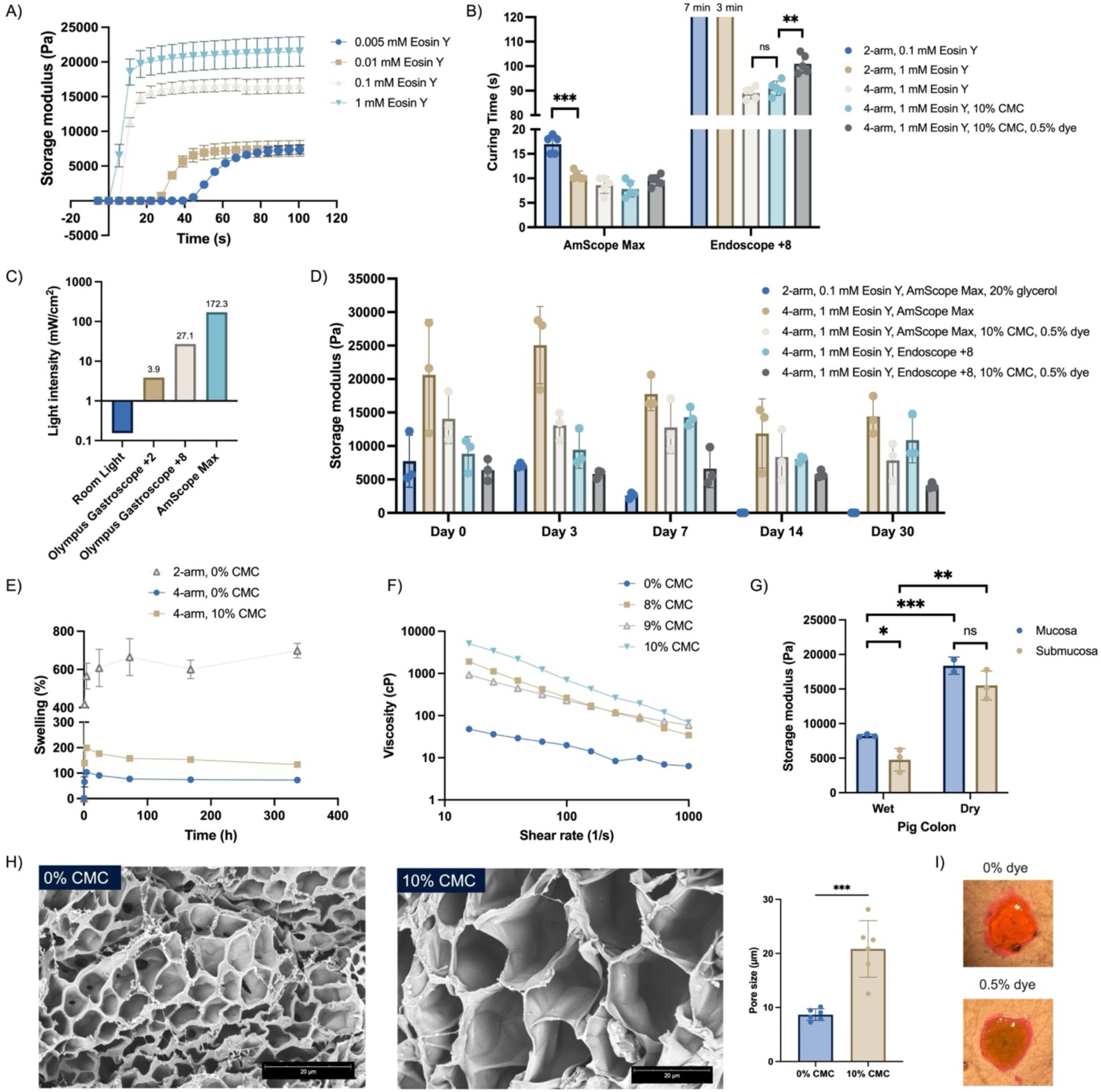
Mechanical and physical characterization of the PolypCure hydrogel. A) In situ photorheometry measuring the storage modulus (G’) and gelation time of 2-arm system hydrogels (n = 5). B) Gelation time of the 2-arm and 4-arm system hydrogels, varying in % CMC, % dye, and the type of light source (n = 5). C) Light intensities of different light sources at 510 nm wavelength. D) Storage modulus (G’) of various hydrogel formulations incubated in PBS at 37 °C over 30 days (n = 3). E) Swelling of the hydrogels incubated in PBS at 37 °C over 14 days (n = 3). F) Viscosities of the 4-arm system hydrogel precursor solutions. G) Storage modulus (G’) of wet and dry pig colon mucosa and submucosa (n = 3). H) SEM images of the 4-arm system hydrogels showing the difference in pore sizes with or without CMC (n = 6). I) Representative images of crosslinked hydrogels with or without dye. Data expressed as mean ± SD; *P < 0.05, **P < 0.01, ***P < 0.001.

Since hydrogel crosslinking is light-dependent,^[38]^ we next examined the spectral flux of four commercial endoscopes: Olympus Gastroscope, Olympus Duodenoscope, Pentax, and Fuji, focusing on the 500-520 nm range that overlaps with the absorbance peak of Eosin Y (∼510 nm).^[53,54]^ While spectral flux and light intensity varied among devices, higher brightness settings and closer proximity consistently yield stronger output (Fig 2C, S5, S6). All endoscopes output substantially lower intensities than AmScope Max light (Fig S6). We selected the Olympus Gastroscope at +8 light settings (hereafter “Endoscope +8”) for its widespread clinical use and sufficient light intensity (27.1 mW/cm^2^ at 510 nm), exceeding the minimum required for effective gelation (Fig 2C).^[38]^

While all formulations cure within 20 s under AmScope Max light, the 2-arm formulations fail to cure within 2 min under the Endoscope +8 light. In contrast, the 4-arm formulations fully gel within 2 min due to higher functionality and improved crosslinking efficiency (Fig 2B). The addition of CMC does not significantly alter gelation time (Fig 2B). Incorporating 0.5% FD&C blue No. 1 dye improves visualization (Fig 2I), a common practice in colonoscopy,^[55]^ without appreciably delaying gelation (Fig 2B, S7). Comparisons between 2- and 4-minute exposures confirm that 2 minutes is sufficient for the gel to reach its stabilized G’ under Endoscope +8 light (Fig S8A). This rapid, one-step gelation (< 2 min) is advantageous compared to hemostatic clips, which require ∼2 minutes per clip, with large lesions typically needing 3-4 clips.^[56–58]^

A key design requirement of the hydrogel adhesive dressing is to achieve mechanical properties compatible with native colon tissue, as mechanical mismatch compromises adhesion or causes premature detachment under dynamic colonic conditions. Rheometry of pig colon tissue reveals that wet mucosa and submucosa possess G’ values of approximately 8 kPa and 5 kPa, respectively (Fig 2G). Thus, an ideal G’ range for the hydrogel is 5-10 kPa. The 2-arm, 0.1 mM Eosin Y formulation cures under AmScope Max light to afford a G’ of ∼6 kPa, consistent with its lower crosslinking density. In contrast, the 4-arm, 1 mM Eosin Y formulations cured under the same light source achieve G’ values exceeding 10 kPa, while curing under Endoscope +8 light produces G’ values between 5-9 kPa (Fig 2D), as expected from the lower light intensity.^[38]^ The addition of CMC and dye slightly reduces G’. Notably, our optimized formulation (4-arm, 1 mM Eosin Y, 10% CMC, 0.5% dye) exhibits G’ values within the desired range, matching the mechanical properties of colon tissue. We also measured the viscosity of the precursor solution to assess its injectability for subsequent catheter-based delivery studies and found it suitable (Fig 2F).

Since delayed bleeding can occur up to 28 days post polypectomy,^[59,60]^ we designed the hydrogel to maintain stability for at least 30 days. Given the colon’s near-neutral pH (6.6-7.5),^[7,61]^ in contrast to the stomach’s acidity (∼pH 2), we evaluated hydrogel stability by incubating samples in PBS at 37 °C for over 30 days and measuring the storage modulus (G’) over 30 days. To enhance *in vivo* durability, we selected amide-linked 4-arm PEG-NB, previously shown to remain stable for 30 days in subcutaneous implants.^[62]^ While the ester-linked 2-arm formulation fully degraded by day 14, the 4-arm hydrogels retain structural integrity throughout the study period (Fig 2D). The optimized formulation maintains G’ > 5 kPa through day 14 and decreases only slightly to ∼ 4 kPa by day 30, confirming its suitability for prolonged colon retention. Considering the colon epithelial turnover rate of 5-7 days under normal conditions and 2-4 days in ulcerative states,^[63–65]^ we expect the hydrogel to detach naturally with the sloughing mucosal layer and be eliminated via feces once re-epithelization is complete.^[26]^

To assess crosslinking density and microstructure, we performed swelling studies and SEM imaging. Hydrogels reach swelling equilibrium after 24 hrs. The 4-arm system exhibits significantly lower swelling (< 200%) compared to the 2-arm formulation (Fig 2E), indicating a denser network. Addition of 10% CMC increases swelling to 176% at 24 hr (Fig 2E). SEM reveals that the average pore size increases from 8.6 µm to 20.8 µm with 10% CMC (Fig 2H, S10). Although chemically inert, CMC may sterically hinder thiol-norbornene crosslinking, leading to increased pore size and swelling. Nevertheless, the pore size of the optimized 4-arm, 10% CMC formulation (∼ 20 µm) remains substantially smaller than that of many hydrogels previously reported for GI applications (> 100 µm).^[26,46,47,66]^ The combination of step-growth polymerization and pre-mixed delivery contributes to the homogeneity and fine microstructure of PolypCure, enhancing its mechanical integrity, adhesive strength, and barrier function.

In summary, the optimized PolypCure formulation (4-arm, 1 mM Eosin Y, 10% CMC, 0.5% dye) gels rapidly within 2 min under standard endoscope light, forming a densely crosslinked network with small pores and moderate swelling. As a result, the hydrogel exhibits mechanical properties compatible with colon tissue and maintains stability in a simulated colonic environment for over 30 days. These findings provide a robust framework for the structure-property relationships of visible-light-crosslinked thiol-norbornene hydrogels and support the evaluation of PolypCure for minimally invasive wound sealing.

### 2.2. Adhesion and delivery of the PolypCure system

Adhesion in the colon environment is particularly challenging due to constant peristaltic motion, the presence of fluids and mucus.^[7,67]^ To assess the adhesive performance of the PolypCure hydrogel, we conducted lap shear tests using *ex vivo* pig colon mucosa and submucosa. We cured the 20 wt% hydrogel *in situ* on tissue surfaces using white light (Fig 3A). Adhesion arises from three likely mechanisms: 1) mechanical interlocking as polymer precursors diffuse into the tissue prior to gelation; 2) covalent bonding between PEG-NB or PEG-SH and tissue thiol groups; and 3) non-covalent interactions including hydrogen bonding and physical entanglement (Fig 1C).

**Figure 3.**
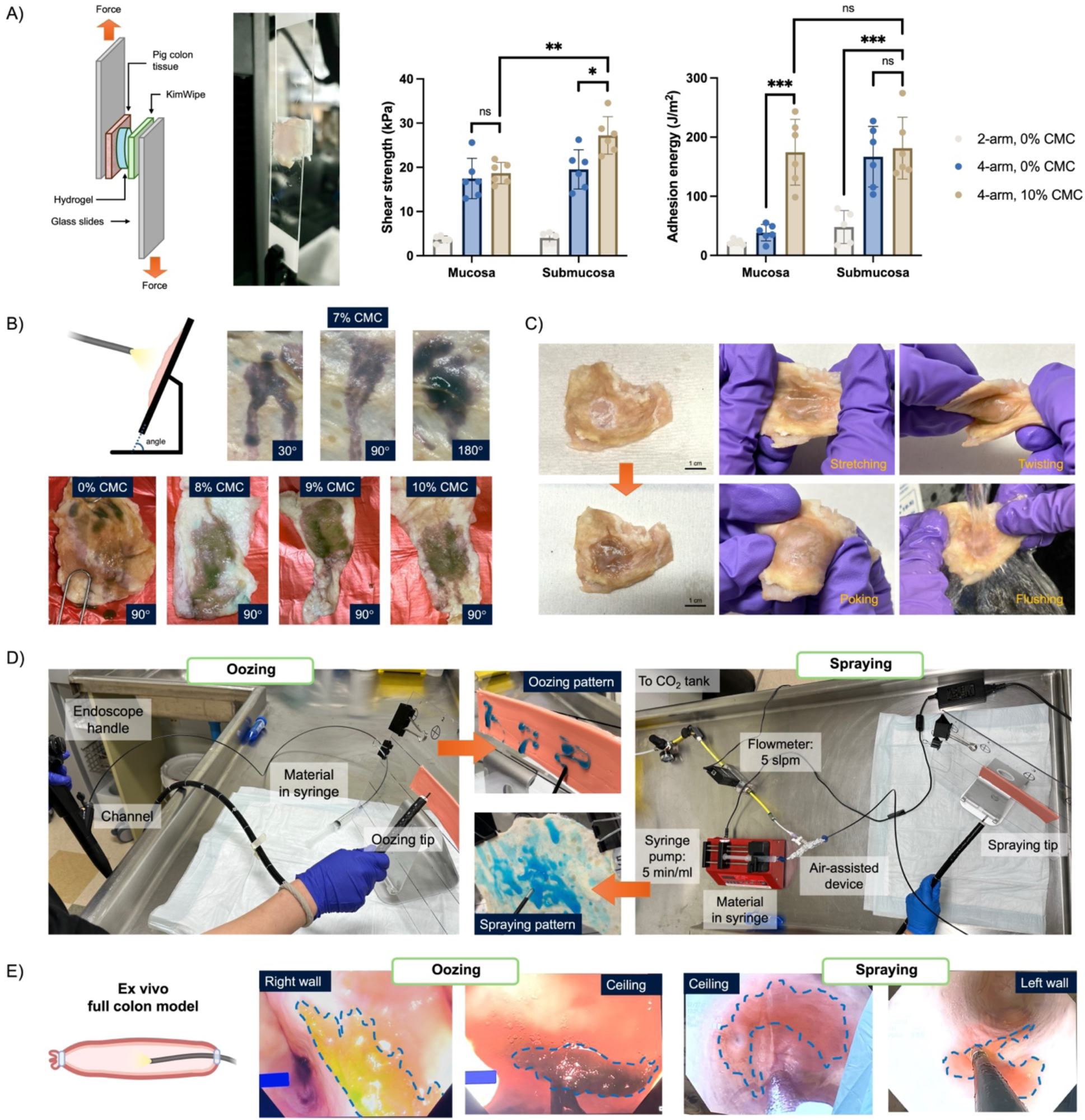
Adhesive properties, delivery methods, and lesion coverage performance of the PolypCure system in *ex vivo* colon models. A) Lap shear strength and adhesion energy of the hydrogels on colon mucosa and submucosa (n = 5 or 6). B) Dripping patterns of the crosslinked 4-arm hydrogels on colon tissue fixed on a slope at varying angles. C) Hydrogels covering colon submucosal lesions exhibit strong tissue adhesion, remaining intact under stretching, twisting, poking and flushing. D) Experimental setup for oozing delivery with a plain black catheter and spraying delivery with the air-assisted device. Oozing and spraying result in distinct material distribution patterns. slpm: standard liters per minute. E) Targeting, curing, and adhesion of the hydrogel at different sites within the *ex vivo* full colon model, delivered via oozing or spraying methods. Data expressed as mean ± SD; *P < 0.05, **P < 0.01, ***P < 0.001.

The 4-arm formulation exhibits significantly higher shear strength (> 18 kPa) than the 2-arm system, surpassing that of commercial adhesives such as Tisseel and Coseal,^[68]^ likely due to tighter crosslinking and reduced pore size. Incorporation of 10% CMC further enhances adhesion strength to ∼27 kPa on submucosa (Fig 1C). Adhesion to submucosa is generally higher than mucosa, likely due to reduced hydration and a denser collagen-rich matrix, which supports both covalent and non-covalent interactions.^[69,70]^ Without CMC, adhesion energy is higher on submucosa. With CMC, adhesion energy on mucosa increases substantially (Fig 3A), as CMC promotes both internal energy dissipation and interfacial hydrogen bonding with the mucus layer. Importantly, the optimized PolypCure formulation robustly adheres to both mucosa and submucosa, and withstood mechanical perturbations such as stretching, twisting, poking, and flushing (Fig 3C).

To ensure effective adhesion across all anatomical orientations of the colon (e.g., floor, wall, ceiling), the hydrogel precursor solution must possess sufficient viscosity to resist gravitational dripping.^[26,71]^ We assessed dripping behavior using an *ex vivo* slope model, where we applied the precursor solution to inclined colon tissue and cured with white light. The 90° incline produces the most significant dripping and, thus, was used as the standard (Fig 3B). To balance anti-drip behavior and catheter injectability, we aimed for a viscosity range of 60–90 cP at a shear rate of 10³ s⁻¹, typical for injection or spraying applications.^[72,73]^ The PEG-based precursor solution without thickeners possesses a low viscosity (< 10 cP) and pronounced dripping (Fig. 2F, 3B). Among the thickeners screened, including glycerol, methylcellulose, gelatin, xanthan gum, and CMC, only CMC sufficiently enhances viscosity and dispersibility (Fig S10). Interestingly, CMC displays greater shear thinning behavior in the presence of PEG: at 2% CMC, viscosity decreases upon PEG addition from 255 cP to below 25 cP (Fig. S8C), likely due to PEG’s plasticizing effect.^[74]^ The 10% CMC formulation achieves a viscosity of 68.9 cP at 10³ s⁻¹, effectively preventing dripping while remaining easily injectable through a standard catheter (OD: 0.079”, ID: 0.060”). The precursor solution stays at the target site until gelation, enabling consistent curing across all colon orientations.

To evaluate clinical usability and optimize application strategies, we tested two delivery modes—oozing and spraying—using an *ex vivo* intact pig colon model. An endoscope was inserted into the colon to simulate *in vivo* procedures and deliver hydrogel to various intraluminal sites (Fig 3E). In the oozing method, the hydrogel precursor solution flows directly through a standard black catheter (Fig 3D). This simple catheter design provides good injectability, smooth passage through the endoscope, minimal risk of clogging, and spreadability of the material. By combining catheter steering with light activation, PolypCure precisely targets lesion sites and cures *in situ* for complete coverage (Fig. 3E). For spraying, we used an air-assisted device developed by Boston Scientific and injected the hydrogel precursor solution at 5 mL/min with a CO_2_ flow rate of 5 slpm to generate a fine mist of droplets (Fig. 3D). Compared to oozing, spraying produces a thinner (< 2 mm), more uniform hydrogel layer with broader coverage and enables a more standardized application protocol (Fig 3E, S11). Notably, both delivery methods achieve precise lesion targeting and mucosal adhesion, withstanding catheter prodding (Mov S1). The cured hydrogel forms a non-adhesive surface, reducing the risk of non-specific adhesion or bowel obstruction.

Together, these results demonstrate that the PolypCure system leverages standard endoscopic tools to enable precise delivery and robust adhesion across all orientations of the *ex vivo* colon. The single-lumen catheter simplifies application, ensures thorough precursor mixing, and supports both direct and aerosolized delivery, underscoring its translational potential.

### 2.3. Biocompatibility, hemostasis and barrier functions of the PolypCure hydrogel

The PolypCure hydrogel is composed primarily of PEG and CMC by weight, both of which are used in FDA-approved products, as well as in pharmaceutical and medical device pre-clinical applications.^[75,76]^ To assess cytocompatibility, we first determined the IC_50_ of the photoinitiator Eosin Y using MTS assays on NIH-3T3 fibroblasts and Caco-2 colon epithelial cells. The IC_50_ values are 1.57 mM and 3.99 mM, respectively (Fig. 4A), both exceeding the 1 mM concentration used in our optimized formulation. Moreover, *in vivo* Eosin Y concentrations are expected to be even lower due to photoinitiator depletion during crosslinking and subsequent diffusion.^[77,78]^ Transwell-based cytotoxicity assays show that all tested formulations maintained >90% cell viability after 24 hr incubation with NIH-3T3 cells (Fig. 4B), confirming the cytocompatibility of the hydrogel. These findings are consistent with prior reports on PEG-based norbornene-thiol hydrogels used in cell encapsulation and tissue engineering applications.^[41,43,79–81]^

**Figure 4.**
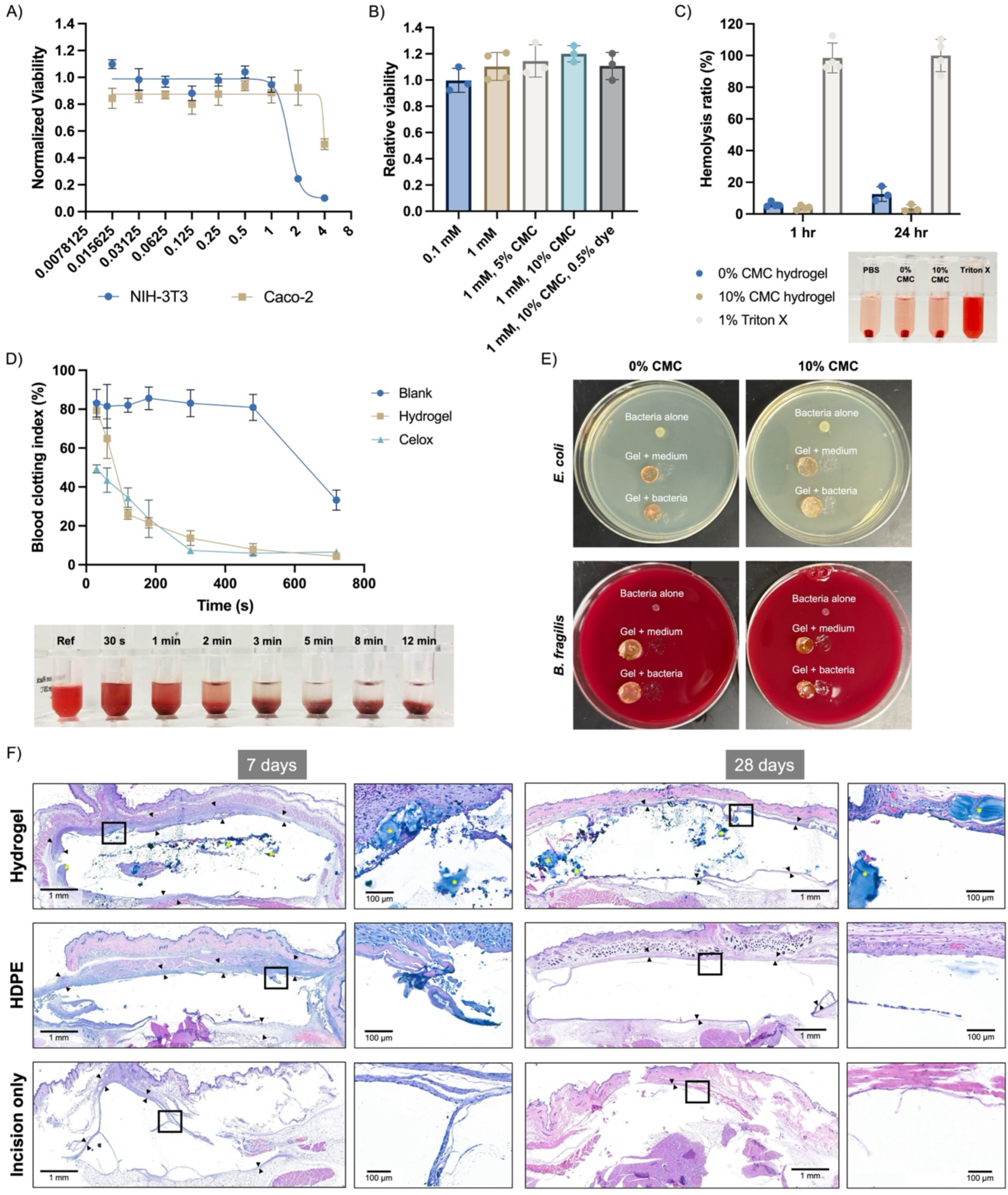
***In vitro* and *in vivo* biocompatibility, hemocompatibility, hemostatic efficacy, and bacteria barrier function of the PolypCure hydrogel system.** A) Cytotoxicity of Eosin Y on NIH-3T3 and Caco-2 cells assessed via MTS assay (n = 6). B) Relative viability of NIH-3T3 cells following 24 hr incubation with hydrogels in a transwell assay (n = 3). C) Hemolysis ratio of hydrogels incubated with red blood cells (RBCs) for 1 or 24 hr (n = 3 or 4). Representative photograph shown at 1 hr. D) Blood clotting index (BCI) of hydrogels compared to blank and Celox powder controls (n = 3). Representative photograph of the hydrogel group. E) Agar plate assays demonstrating that the hydrogel blocks migration of *E. coli* and *B. fragilis* during overnight incubation (n = 3). F) H&E staining of local tissue from the mouse subcutaneous implantation study (n = 4 per group per time point). Yellow asterisks mark residual hydrogel; black arrowheads indicate the fibrous capsule surrounding the implant. Most implants were removed or lost during processing. Data expressed as mean ± SD.

We next evaluated hemocompatibility using a hemolysis assay.^[82]^ At 1 hr, the supernatant from the hydrogel groups appears similar to the PBS control, in contrast to the bright red of the Triton X positive control (Fig 4C). The 0% CMC hydrogel shows hemolysis ratios of 5.90% at 1 hr and 12.57% at 24 hr, while the 10% CMC formulation reduces hemolysis to 3.62% and 3.89%, respectively (Fig. 4C), remaining below the acceptable threshold (< 5%) for hemostatic biomaterials.^[83,84]^ This reduction is due to the hydrophilic and anionic nature of CMC, which repels negatively charged red blood cell membranes.^[85]^

The PolypCure system is designed to provide immediate hemostasis post-polypectomy by forming a physical barrier and initiating clotting via contact activation of the coagulation cascade.^[86]^ Thus, we evaluated clotting efficacy using the blood clotting index (BCI), where values below 20% indicate excellent hemostatic properties.^[87]^ In untreated activated rat whole blood, spontaneous clotting occurs at 12 min with a BCI of 33%. In contrast, the PolypCure hydrogel reduces BCI from 79.3% at 30 s to below 25% by 2 min (Fig 4D), demonstrating rapid clot formation concurrent with gelation. Clotting is visually apparent before 30 s and completed by 3 min. As a positive control, Celox hemostatic granules (polymer-mass equivalent) achieve comparable clotting kinetics. Unlike hemostatic clips, which often fail to ensure hemostasis and risk secondary tissue damage,^[88]^ the PolypCure hydrogel forms a conformal matrix at the bleeding site to achieve reliable and atraumatic hemostasis.

Following colon polypectomy, the compromised intestinal barrier increases susceptibility to bacterial infiltration. Thus, we assessed the barrier function of the PolypCure hydrogel using an agar-based assay against *E. coli* and *B. fragilis*, two abundant gut bacteria implicated in impaired healing and postoperative infections.^[13,89,90]^ We applied bacteria (OD_600_= 1.0) to the apical surface of 2 mm-thick hydrogel discs. After 24 hr, no bacterial growth is present beneath either 0% or 10% CMC hydrogels (Fig 4E), indicating effective blockage of microbial migration. We confirmed bacterial viability by flipping the hydrogel and re-incubating (Fig S12). Although bacterial dimensions are smaller than the individual pores (< 30 µm),^[13]^ the hydrogel’s dense, tortuous network impedes bacterial translocation, thus serving as a physical barrier to protect the lesion from bacterial infection.

Next, we assessed *in vivo* biocompatibility in a mouse subcutaneous implantation model. We implanted the PolypCure hydrogel into the subcutaneous space on the back of the animals (Fig S13), with USP high-density polyethylene (HDPE) implants as a positive control and an incision-only group as the incision-only procedure control. We observed no abnormal mouse behaviors or weight loss (Fig. S14B), and white blood cell counts remain comparable across all groups on both days 7 and 28 endpoints (Fig. S14A). H&E staining of major organs (heart, lung, liver, spleen, kidney) show no pathological abnormalities (Fig. S15), indicating the hydrogel does not elicit systemic toxicity.

Local tissue histology confirms implant retention at day 28, although most hydrogel materials were lost during H&E processing, leaving white spaces in the subcutaneous pocket (Fig 4F). Occasional residual hydrogel (yellow asterisks) is visible. At day 7, loosely organized fibroblasts, neutrophils, and macrophages are present around the implants, consistent with a mild chronic inflammatory response. By day 28, a compact fibrous capsule (black arrowheads) forms with reduced immune cell infiltration. Capsule thickness and cellular composition are comparable between PolypCure and HDPE implants, suggesting a similar level of foreign body response. The incision-only group also exhibits mild fibrosis, likely due to surgical trauma. These findings align with the expected host response to implanted biomaterials and are consistent with prior studies on PEG-based norbornene-thiol hydrogels.^[62,91–93]^ Overall, the mice tolerate the PolypCure hydrogel, supporting good *in vivo* biocompatibility.

Collectively, these results confirm the *in vitro* and *in vivo* biocompatibility and hemocompatibility of the PolypCure hydrogel, suggesting its safety for further *in vivo* evaluation. It promotes rapid hemostasis and prevents bacterial migration, provides mechanical protection to reduce bleeding, and inhibits postoperative infection.

### 2.4. *In vivo* evaluation of the PolypCure system in a pig EMR model

We evaluated the *in vivo* performance of the PolypCure system using a Yorkshire pig colon endoscopic mucosal resection (EMR) model, which recapitulates the anatomical structure, mucosal composition, and tissue repair processes of the human colon.^[94–96]^ This model also accommodates standard endoscopic instruments, enabling clinically relevant procedures. We created EMR lesions (∼20–30 mm in diameter) by submucosal saline injection followed by mucosal excision with a Captivator II snare (Boston Scientific). Untreated lesions served as negative controls and were formed at the proximal end to minimize cross-contamination. We used an Olympus gastroscope for access and visualization.

Achieving *in vivo* adhesion in the colon is particularly challenging due to continuous peristalsis and a highly hydrated mucosal surface.^[7,26,67]^ We first validated the feasibility of PolypCure delivery and adhesion *in vivo*. For oozing delivery, we extruded 1 mL of hydrogel precursor solution through a standard black catheter onto a right wall lesion. The solution remains in place due to its sufficient viscosity and cures under Endoscopic +8 light within 1–2 minutes (Fig 5B). The reflective mucosal surface and elevated body temperature likely further accelerated gelation. Application occurs smoothly, with no catheter clogging or obstruction of endoscopic view (Mov S2). The hydrogel remains tightly adhered to the lesion for over 90 min and resists aggressive prodding with an interject needle (Mov S2), indicating strong adhesion capable of withstanding colon motility. Necropsy confirms complete hydrogel coverage of the lesion (Fig 5B). Residual material (black arrows) remains on the submucosa even after histological processing (Fig 5C), confirming robust adhesion.

**Figure 5.**
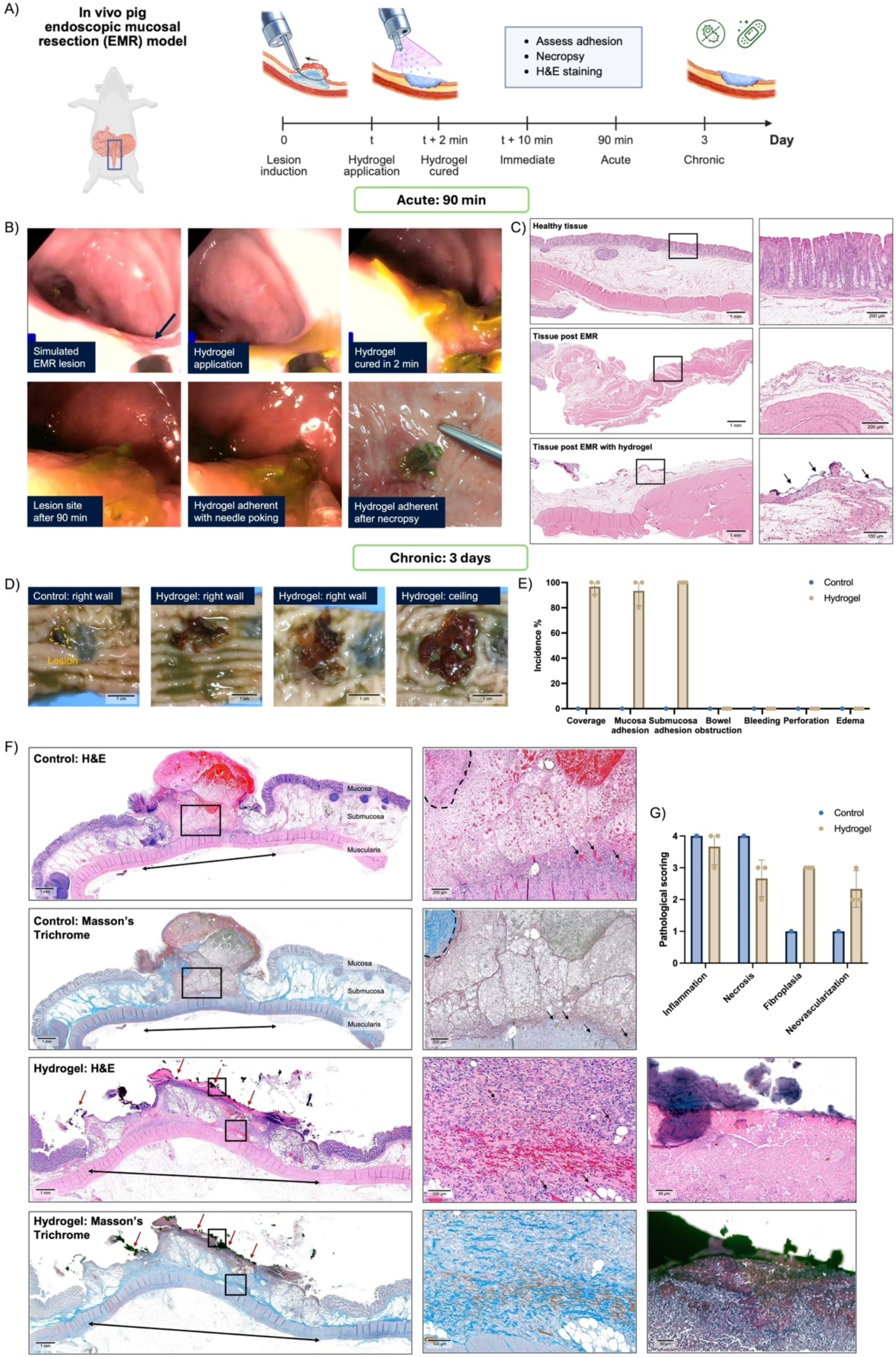
***In vivo* evaluation of the application, adhesion, and wound healing performance of the PolypCure system in a pig endoscopic mucosal resection (EMR) model.** A) Schematic of the experimental timeline. Following EMR lesion creation, hydrogel was applied and cured in situ. Adhesion was assessed endoscopically at 10 min and 90 min. Animals were euthanized at 90 min or on day 3 for necropsy and histopathological analysis. B) Endoscopic images showing oozing application of the hydrogel solution to an EMR lesion. The hydrogel cured within 2 min under endoscope light and remained adhered to the lesion at 90 min *in vivo* and at necropsy. C) H&E staining of healthy tissue, post-resection tissue, and hydrogel-treated tissue at 90 min. Black arrows indicate residual hydrogel on colon submucosa; most hydrogel was lost during histological processing. D) Gross necropsy images of control and hydrogel-treated sites at day 3. E) Quantification of hydrogel coverage, adhesion to mucosa and submucosa, and any observed adverse effects. F) H&E and Masson’s trichrome staining of lesion sites at day 3. Lesion size (double-headed arrows), hydrogel presence (red arrows), fibroplasia (dashed lines), and neovascularization (black arrows) are indicated. High-magnification images confirm hydrogel retention and adhesion to the colon submucosa. G) Histopathological scores of lesion sites at day 3. Data expressed as mean ± SD.

We also evaluated spraying delivery on a pig stomach EMR lesion. An air-assisted device atomizes the hydrogel precursor solution and forms a thin, uniform coating over the lesion within 30 s (Fig S16, Mov S3). After 2 min of light exposure, the hydrogel adheres to the stomach tissue and resisted mechanical disruption, especially at the submucosa. Forced detachment by mechanical manipulation with the scope causes tissue damage and bleeding, highlighting robust interfacial integration and effective adhesion even in the stomach. However, spraying delivery presents practical challenges. During colonoscopy, the lumen is already insufflated with CO_2_.^[97]^ Additional gas flow from spraying may elevate intraluminal pressure, requiring careful pressure control (10–25 mmHg) to avoid mucosal trauma and maintain spray pattern.^[98–100]^ This necessitates additional equipment (e.g., flow meters, pressure sensors) and operator training, potentially hindering clinical adoption by increasing the cost and complexity of the procedure. In contrast, oozing delivery requires only a syringe and catheter, offering a simple and low-cost delivery method using standard off-the-shelf components. While many spontaneously gelling multi-component hydrogels rely on spraying for *in situ* mixing,^[13,26,45,46,66]^ our white light-activated *in situ* crosslinking approach enables precursor pre-mixing and simple oozing delivery, thus facilitating clinical translation. Accordingly, we adopted oozing delivery for subsequent studies.

To assess longer term adhesion and potential healing effects, we conducted a 3-day chronic study in the pig colon EMR model. We created four lesions; one was left untreated, while three receive the PolypCure hydrogel (Fig S17). We applied the hydrogel precursor solution under low-intensity Endoscopic −8 light to prevent premature curing and cured *in situ* under Endoscopic +8 light for 2 min (Mov S4). For difficult-to-access sites, one can incrementally ooze the hydrogel to ensure complete lesion coverage. At day 3 necropsy, all treated lesions remain fully covered by thin, flat hydrogel layers with strong adhesion (Fig. 5D). Mechanical rubbing and scraping reveals stronger submucosal than mucosal adhesion, consistent with prior *in vitro* findings. No signs of bleeding, perforation, bowel obstruction, or edema are present (Fig 5E), confirming *in vivo* safety. These results highlight the hydrogel’s ability to maintain stable adhesion in the dynamic colon environment for at least 3 days, potentially reducing delayed bleeding and perforation, typically observed after post-polypectomy.^[18,24,25,101]^

We performed H&E and Masson’s trichrome staining on the lesion sites and scored for inflammation, necrosis, fibroplasia, and neovascularization (Fig 5F, Fig 5G, Fig S19). The untreated lesion exhibits protruding ulceration with extensive necrosis and inflammatory cell infiltration at the lesion margins (Fig 5F). Fibroplasia (dashed lines) and neovascularization (black arrows) are few, consistent with acute inflammation phases of early wound healing.^[96,102]^ In contrast, PolypCure-treated lesions show physical containment of the ulcer, and signs of accelerated tissue repair (Fig 5F, Fig S18). Inflammatory cell infiltration reduces, due to the hydrogel’s barrier function that shields the wound from microbial invasion and mechanical irritation. While necrosis persists in the mucosa and submucosa, granulation tissue formation, submucosal reconstitution, and mucosal re-epithelialization are present. Fibroplasia and neovascularization (black arrows) appear more extensive and organized throughout the submucosa, suggesting more controlled tissue remodeling.^[96,103]^ High-magnification images further confirm hydrogel retention (red arrows) and strong adhesion to the submucosa. Although the sample size was insufficient for statistical significance, hydrogel-treated lesions show a consistent trend toward reduced inflammation and necrosis, along with enhanced fibroplasia, neovascularization, and re-epithelialization (Fig. 5G). These findings suggest that the PolypCure hydrogel physically isolates the wound to create a more sterile and less inflammatory wound bed, thus accelerating natural wound healing.

Together, these results demonstrate that the PolypCure hydrogel effectively seals large EMR lesions using standard endoscopic tools, maintains stable *in vivo* adhesion for at least 3 days, and promotes wound protection and healing. These outcomes support its potential to reduce post-polypectomy complications, including intraprocedural bleeding, delayed bleeding, and delayed perforation, although longer-term studies (up to 28 days) are needed to fully validate therapeutic efficacy.

## 3. Outlook

Colon polypectomy is a common preventive procedure for colorectal cancer, performed in approximately half of individuals over 50 years of age.^[104,105]^ However, polyp removal via EMR or ESD creates exposed wounds that are susceptible to bleeding, perforation, and infection.^[7,11,28]^ Mechanical clips are currently the standard of care for managing these wounds, but they are technically demanding, costly, time-consuming, and can cause secondary trauma.^[7,13,14,17]^ Moreover, closing large lesions (> 2 cm in diameter) often requires multiple clips and may be impossible due to poor accessibility.^[7,17,58,106,107]^

The concept of shielding post-resection colon wounds with biomaterials was introduced in 2014 using polyglycolic acid sheets and fibrin glue.^[108,109]^ Since then, various hemostatic powders and surgical sealants have been explored but have failed to reach the clinic due to inadequate adhesion or complex application.^[13,15,17,110]^ More recently, injectable hydrogels are being developed specifically to seal colon lesions and provide mechanical protection. Crosslinking strategies include N-hydroxysuccinimide (NHS) ester-amine, maleimide-thiol, Schiff base, radical polymerization, and thermoresponsive systems.^[7,13,26,46,47,111–113]^ Among these, a polyethyleneimine (PEI)-modified Pluronic micelles and oxidized dextran formulation demonstrates one of the longest retention times *in vivo* (3–7 days),^[26]^ while others provide varying degrees of wound protection and healing enhancement.^[26,47,66,111,112]^

Despite these advances, existing hydrogel formulations for colon applications rely exclusively on spontaneous *in situ* crosslinking, which presents significant limitations in usability and performance. Slow-gelling systems require pre-polymerization prior to application,^[66,105]^ increasing the risk of premature gelation and catheter clogging. Fast-gelling systems necessitate dual- or tri-lumen catheters to co-deliver separate components to the lesion or involve multi-step spraying protocols.^[13,26,45–47]^ These delivery approaches are operationally complex and lead to heterogeneous mixing on tissue, which compromises crosslinking efficiency, mechanical robustness, and adhesive strength.

In this work, we present the first hydrogel system that leverages the white light source embedded in standard endoscopic instruments to achieve spatiotemporally controlled *in situ* crosslinking within the GI tract. The PolypCure hydrogel consists of a pre-mixed precursor solution containing 4-arm PEG-NB, 4-arm PEG-SH, CMC, and the photoinitiator Eosin Y. Upon oozing delivery, the precursor solution remains localized at the lesion site via CMC-mediated viscosity and non-covalent interactions. Endoscopic white light initiates rapid hydrogel crosslinking within 2 min, forming strong tissue adhesion through covalent bonding and mechanical interlocking. This design supports universal application to colon lesions of various sizes and locations, enables rapid hemostasis and stable adhesion, and is fully compatible with standard instruments. The resulting hydrogel exhibits a highly uniform structure with small pore size (∼20 µm), low swelling ratio (< 200%), appropriate mechanical properties (G’ ∼ 6 kPa), high shear strength (>15 kPa) and strong adhesion energy (>150 J/m^2^). It remains stable in simulated colon conditions for up to 30 days and effectively prevents bacterial penetration, fulfilling key design criteria for a colon-specific wound dressing.

While thiol–norbornene hydrogels are extensively explored for *in vitro* cell culture and bioprinting,^[41,114]^ their *in vivo* use is limited.^[62,115,116]^ This study presents the first report of a photocrosslinked norbornene–thiol hydrogel adhesive dressing applied *in vivo* via a minimally invasive approach. We demonstrate *in vitro* cytocompatibility and hemocompatibility, and confirm *in vivo* biocompatibility via subcutaneous implantation studies, using FDA-approved HDPE as a benchmark control.

Most notably, the PolypCure hydrogel is efficacious in a pig colon EMR model with clinically challenging lesions (> 2 cm in diameter, located on the wall or ceiling). The hydrogel remains fully adhered for at least 3 days under harsh *in vivo* conditions, characterized by high mucus content, bacteria load, and constant mechanical motion.^[7,26]^ Histological analysis reveals reduced inflammation and necrosis along with enhanced fibroplasia and neovascularization by day 3. The data suggest that the hydrogel accelerates re-epithelialization and tissue remodeling.^[96]^ These results highlight the hydrogel’s potential to reduce delayed bleeding and perforation, positioning PolypCure as a strong candidate for clinical translation in post-polypectomy wound management. Further, given that polymerization occurs at white light intensities as low as 3.6 mW/cm^2^,^[38]^ we anticipate broad compatibility across clinical platforms, enabling potential use in other minimally invasive procedures such as upper GI endoscopy, laparoscopy, or arthroscopy.^[117,118]^

## CREDIT authorship contribution statement

**Xingyu Hu:** Conceptualization, Methodology, Formal analysis, Investigation, Data Curation, Writing – Original Draft, Writing – Review & Editing, Visualization, Project administration. **Rong Liu, George A. O’Toole, Shawna Pratt, Samantha Berry, Michal Jones, Aaron Rosenthal:** Methodology, Formal analysis, Investigation, Data Curation. **Yin P. Hung**: Investigation, Formal analysis. **Carolina Villarreal:** Conceptualization, Resources, Writing – Review & Editing, Supervision. **Mark W. Grinstaff:** Conceptualization, Resources, Writing – Original Draft, Writing – Review & Editing, Supervision, Funding acquisition.

## Acknowledgements

The authors would like acknowledge Allison Kumarasena and Kevin Woods from Boston Scientific for their valuable insights and assistance with the *in vivo* animal model experiments. The content is solely the responsibility of the authors and does not necessarily represent the official views of the National Institute of Health.

## Declaration of Generative AI and AI-assisted Technologies in the Writing Process

During the preparation of this work the author(s) used no tools or services.

## 4. Experimental Section

***Polymer synthesis*:** 4-arm PEG-norbornene (4-arm PEG-aNB, MW 5000) was synthesized according to an established protocol with slight modification.^[48,49]^ Briefly, 5-norbornene-2-carboxylic acid (5.52 g, 40 mmol, Sigma Aldrich) was reacted with N,N’-dicyclohexylcarbodiimide (4.12 g, 20 mmol, Sigma Aldrich) in dichloromethane (DCM) for 2 hr to form norbornene anhydride and by-product dicyclohexylurea. Norbornene anhydride was filtered through a fritted funnel and added into a second flask containing pre-dissolved 4-arm PEG-amine (10 g, 2 mmol, MW 5000, JenKem Technology USA), 4-(dimethylamino)pyridine (97.6 mg, 0.8 mmol), and N,N-diisopropylethylamine (1.742 ml, 10 mmol) in DCM. The reaction was allowed to proceed overnight at room temperature. The crude was washed with citric acid, sodium bicarbonate, and brine, then precipitated in cold diethyl ether twice, and dried under vacuum for 3 days (white solid, 67% yield). The degree of functionalization (> 95%) was characterized by proton NMR.

2-arm PEG-thiol (2-arm PEG-eSH, MW 2000) was synthesized via functionalizing PEG with dithiol using 3-mercaptopropionic acid (MPA).^[50]^ PEG (4 g, 2 mmol, 2000 MW) and MPA (2.12 g, 20 mmol) were dissolved in toluene and heated to 50 °C before p-toluenesulfonic acid (38 mg, 0.2 mmol) was added to the mixture. The solution was further heated to 140 °C, stirred, and refluxed overnight. The crude was purified by precipitating in cold diethyl ether twice and dried under vacuum for 3 days (white solid, 92% yield). The degree of functionalization (> 84%) was characterized by proton NMR. 4-arm PEG-thiol (4-arm PEG-mSH, MW 5000) was purchased from JenKem Technology USA and used as received.

***Hydrogel preparation*:** Hydrogel samples were synthesized at various Eosin Y, carboxymethyl cellulose (CMC, 1500-3000 cP, Sigma Aldrich), and blue dye (FD&C Blue No. 1) concentration with a ratio of norbornene (NB) to thiol (SH) of 1:1 and a total polymer concentration of 20 wt%. Specifically, 4-arm PEG-aNB, 4-arm PEG-mSH or 2-arm PEG-eSH, and the blue dye were first dissolved in PBS and sterile filtered through 0.22 µm PES filters. UV-sterilized CMC or other thickeners (glycerol, methylcellulose 4000 cP, methylcellulose 1500 cP) was then slowly added to the vial and vortexed to mix. Next, sterile filtered Eosin Y was added to the mixture. The polymer solution was loaded into foil-wrapped syringes and kept in the dark until further uses. Various white light sources were used to cure the solution approximately 2 cm away into custom-designed cylindrical PTFE molds (8 mm or 12 mm in diameter). Gelation time was determined by the inversion test (tilting or inverting every 5 s until the gel remained entirely at the bottom).^[119]^

***Rheological characterization*:** Rheological measurements were conducted using a TA Instruments DHR-2 rheometer following previously published protocols.^[120]^ For pig colon mucosa and submucosa tissues, oscillatory strain sweeps were performed from 0.01% to 10% at 5 rad/s at 37 °C using 8 mm plates. Storage modulus (G’) was reported as an average across the strain sweep before it dropped by more than 5%. Dry pig colon tissue was obtained by allowing fresh tissue to dry at room temperature for 1 hr. For hydrogel samples, frequency sweeps were performed at 1% strain (within the linear viscoelastic region) from 0.1 to 100 rad/s at 37 °C using 8 mm plates. Storage modulus (G’) and loss modulus (G”) of each sample was reported as an average across the frequency sweep before it dropped by more than 5%. In situ photorheometry was performed using AmScope white light at max settings (150W, HL150-A). 10 µl polymer solution containing 4-arm PEG-aNB and 2-arm PEG-eSH was pipetted onto the middle of the 8 mm transparent parallel plate connected to the white light. Time sweep was conducted at 1% strain and 5 rad/s frequency for each hydrogel (n = 5) at a gap of 280 µm for 100 s after white light was turned on. Gelation point was defined as the time when G’ surpasses G”. The viscosity of the hydrogel precursor solution was measured using the 20 mm 2° cone geometry at a shear rate of 10/s to 1000/s at 37 °C. 45 µl solution was loaded onto the plate and the truncation distance was set to 70 µm.

***Swelling, SEM, and degradation*:** Swelling ratio was characterized by submerging and weighing the hydrogel in 3 ml of PBS at 37 °C over 14 days. The hydrogel was weighed at each timepoint. Swelling ratio was calculated using the following formula: Swelling (%) = (W_t_ – W_0_)/W_0_, where W_t_ is the wet mass of the hydrogel at time t and W_0_ is the initial wet mass of the hydrogel following fabrication. Scanning electron microscopy (SEM) images were acquired using a Phenom PROX Desktop SEM at 5 kV, with a working distance of 9-12 mm and an aperture size of 30 µm. Hydrogels were swollen in water overnight, lyophilized, and sputter coated with AuPd for 20 s. Pore area was quantified and analyzed using the built-in threshold and particle analysis tools in ImageJ (v1.54f) following an established protocol.^[121]^ For the degradation test, pre-made hydrogels (n = 3) were submerged in PBS solutions and incubated at 37 °C over 30 days. At each timepoint, we performed a frequency sweep for each sample and reported its storage modulus (G’).

***Light characterization*:** White light intensity (mW/cm^2^) was measured using ThorLab PM100D with S121C sensor at 510 nm wavelength and 2 cm distance. Spectral radiant flux (W/nm) was measured using Gigahertz-Optik BTS256-LED tester at wavelengths between 360 nm and 830 nm. The tested light sources included the AmScope white light (150W, HL150-A), Olympus gastroscope (GIF-2TH180), Olympus duodenoscope (TJF-Q190V), Pentax colonoscope (EC38-i10cL), and Fujifilm colonoscope (EC760R-V/L). Spectral radiant flux was measured for Olympus gastroscope at +1, +2, +4, +6, and +8 settings, while all other devices were measured at +1. Multiple AmScope white light settings were tested to match the Olympus gastroscope for *in vitro* experiments.

***Tissue adhesion studies*:** Fresh moisturized pig colon tissue was used for tissue adhesion studies. The submucosa layer was obtained by scraping the mucosa layer with a scalpel. Lap shear adhesion tests were performed following a modified ASTM F2255 standard “Strength Properties of Tissue Adhesives in Lap-Shear by Tension Loading” using an Intron 5944 Microtester with 100 N load cells.^[119,122]^ Pig colon tissue and Kimwipes were cut into 2.5 cm x 1 cm squares and firmly glued to the glass slides. 100 µl of hydrogel precursor solution was spread onto the tissue, covered by the Kimwipe on other glass slide, and cured by AmScope white light for 2 min. After stabilizing at room temperature for 5 min, the hydrogel and the tissue were pulled apart at a rate of 5 mm/min until a break in adhesion was detected. Shear strength (kPa) was calculated as maximum load F_max_ divided by sample area. Adhesion energy (J/m^2^) was calculated as the area under the force-displacement curve divided by sample area. For the *ex vivo* lesion adhesion study, 2 cm submucosal lesions were created using razor blades (n = 6). 150 µl of hydrogel solution was applied and cured to fully cover each lesion. The material’s adhesive properties were examined through stretching, twisting, poking, and flushing for more than 30 s.

***Catheter designs*:** Multiple in-house catheter designs were developed to optimize the delivery of the hydrogel precursor solution through the standard Olympus gastroscope (GIF-2TH180), with catheter diameters ranging from 0.15 to 0.2 cm. A 9% carbon black cover was incorporated to prevent premature light-induced curing. Various catheter tip designs were evaluated for injectability, endoscope passability (kink resistance), susceptibility to tip clogging, and ability to spread material on tissue. An air-assisted miniature nozzle catheter (Boston Scientific) was designed to generate fine dispersions of fluid droplets. Material delivery was regulated via a syringe pump, while air was supplied from a CO_2_ tank. The syringe pump injection rate and air flow rate (monitored via a flowmeter) were adjusted to achieve optimal spray performance.

***Ex vivo slope tests*:** A slope apparatus was constructed with an adjustable angle ranging from 0° to 180° to simulate the anatomy of the colon. Fresh colon tissue was secured onto the slope for testing. 200 µl of hydrogel precursor solution (4-arm PEG-aNB, 4-arm PEG-mSH, 20 wt%, 1:1 NB:SH, 0.5% dye) was delivered through a plain black catheter onto the colon tissue under Olympus Colonoscope +2 light. The hydrogel was then cured with Olympus Colonoscope +8 light for 2 min, and the dripping and curing patterns were recorded.

***Ex vivo full colon model*:** Fresh, unsectioned colon tissue (> 80 cm in length) was used to create a full colon model. One end of the colon was tied off, and the other end was insufflated with CO_2_ to allow endoscopic insertion. The best performing hydrogel precursor solution (4-arm PEG-aNB, 4-arm PEG-mSH, 20 wt%, 1:1 NB:SH, 10% CMC, 0.5% dye) was applied to various locations within the colon, including the ceiling, right wall, left wall, and floor. For oozing delivery, the plain black catheter dispensed material under endoscope −8 light for 30 s, followed by curing under endoscope +8 light for 2 min. For spraying delivery, the effervescent device was operated at 5 ml/min injection rate and 14 psi CO_2_ pressure. Material was applied under endoscope +2 light for 30 s and cured at endoscope +8 light for 2 min. The endoscope tip was used to prod the hydrogel dressing to assess adhesion. Potential contamination of the catheter tips was monitored and documented.

***In vitro cytotoxicity*:** *In vitro* cytotoxicity studies were performed for the photoinitiator Eosin Y on NIH-3T3 murine fibroblast cells and Caco-2 human colon epithelial cells following our previously published studies.^[123,124]^ Cells were seeded at 5,000 cells per well in 96-well plates and allowed to adhere for 1 day and 2 days for NIH-3T3 and Caco-2 respectively, followed by treatment with photoinitiators for 24 hr. Cell viability and IC_50_ was determined using the MTS assay (n = 6). Blank wells and wells with media were included for background correction. *In vitro* cytotoxicity of hydrogels against NIH-3T3 cells was investigated using 3 µm pore polycarbonate transwell inserts in 12-well plates. Sterile filtered hydrogel precursor solution was cured inside the transwell and incubated with cells for 24 hr. Cell viability was assessed using the MTS assay.

***Hemolysis and blood clotting studies*:** Hemolytic activity of the hydrogels was evaluated following published protocols.^[27]^ Red blood cells (RBCs) were isolated from citrated rat whole blood (Innovative Research) and triple-washed with PBS by centrifugation at 5000 rpm for 10 min. RBCs were diluted to 5% v/v in PBS and aliquoted as per 1000 µl in tubes. 50 µl of hydrogel was added to each tube (n = 4) and incubated at 37 °C water bath for 1 hr or 24 hr. PBS and 1% Triton X were used as negative control and positive control, respectively. The samples were then centrifuged at 5000 rpm for 10 min. Hydrogel samples were removed before centrifugation. The absorbance of the supernatant was measured at 540 nm and the hemolysis ratio was calculated as (OD_sample_ – OD_negative_)/(OD_positive_ – OD_negative_) x 100%.

The blood clotting index (BCI) was measured adapting from a published protocol.^[125]^ Activated blood was prepared by mixing citrated rat whole blood (Innovative Research) with 0.1 M calcium chloride solution at a 9:1 volume ratio. For the hydrogel group, 10 µl of activated blood was added into 50 µl of hydrogel solution and cured under +8 light. A weight-equivalent amount of Celox hemostatic granules (MedTrade Products Ltd.) served as a positive control. The negative control consisted of 10 µl of activated blood alone, allowing natural coagulation. At selected time points, 500 µl of deionized water was added to each tube to release the uncoagulated blood components. The supernatant was collected and absorbance measured at 540 nm. The reference was 10 µl of citrated whole blood in 500 µl deionized water. BCI was calculated as (OD_sample_ / OD_reference_) x 100%.

***Bacteria migration assays*:** Bacteria migration experiments were performed following our previously published protocol.^[119]^ *Escherichia coli* (TL139H) was cultured aerobically overnight on LB agar, while *Bacteroides fragilis* (HAP130N-2B) was cultured anaerobically for 48 hr on blood agar (tryptic soy agar with 5% sheep blood) using the GasPak system. Colonies were harvested into 1 ml PBS, homogenized, centrifuged at 16,000 x *g* for 30 s, resuspended in PBS, and normalized to an OD600 of ∼1.0.

For the assay, hydrogel samples (12 mm diameter x 2 mm height) were placed on agar plates, and 5 µl of bacteria suspension, or the medium only control, was added onto the top of each hydrogel (n = 3). “Bacteria-only” indicates that 5 µl of bacteria suspension was spotted directly onto the agar plate and served as a positive control. Agar plates were incubated at 37 °C for 24 hr either in room air (*E. coli*) or under anoxic conditions (*B*. *fragilis*). To assess bacterial migration, hydrogels were gently lifted to inspect for bacterial growth on the agar plate underneath. Hydrogels were then flipped and incubated for another 24 hr, under the appropriate condition for each organism, to confirm bacterial viability.

***Biocompatibility and safety studies in subcutaneous implants in mice*:** Animal experiments were approved by the Institutional Animal Care and Use Committee of Massachusetts General Hospital (MGH) and conducted in strict compliance with all federal and institutional guidelines. Equal numbers of male and female C57BL/6J mice (6-8 weeks old) were obtained from Jackson Laboratories (Bar Harbor, Maine). Mice were anesthetized with isoflurane inhalation and received a dose of long-acting analgesic (Ethiqa XR 3.25 mg/kg, subcutaneously). Under sterile conditions, a subcutaneous pocket approximately 1 cm wide was created by blunt dissection of the connective tissue on the dorsal side between the shoulders. Either the best performing hydrogel (4-arm PEG-aNB, 4-arm PEG-mSH, 20 wt%, 1:1 NB:SH, 10% CMC, 0.5% dye) or USP high density polyethylene (Sigma Aldrich) was cut into 1 cm x 1 cm squares and subcutaneously implanted in the back of male or female C57BL/6 mice (n = 4 per group per time point). Additional mice that received only incision and blunt dissection served as surgical intervention-only controls. The incisions were closed with rodent wound clips (Reflex 7mm, Roboz Surgical Instrument), which were removed 10-14 days post-surgery. Body weight was monitored on days 1-5, 7, 14, and 28. On day 7 and day 28, mice were euthanized, and blood was collected for white blood cell counts using the Medix Leukotic Bluplus test kit (Fisher Scientific). Tissue surrounding the implant site was harvested and processed for Hematoxylin-Eosin (H&E) staining at the MGH Histopathology Research Core. Histopathological evaluations were performed in a blinded manner by an MGH attending pathologist. Degrees of inflammation, fibrosis, necrosis, and immune responses were assessed on cross-sections of the individual implants and the surrounding tissues. Major organs (liver, kidney, spleen, lung, and heart) were also collected and H&E stained to assess systemic toxicity.

***In vivo pig acute EMR model*:** The pig studies were performed at CBSET (Lexington, MA), an AAALAC International-accredited facility, in compliance with all relevant regulations for laboratory animal care and use. The animal procedures were performed under the approved IACUC protocols. A simulated endoscopic mucosal resection (EMR) lesion was created by first injecting a lifting agent (1% indigo carmine in saline) into the submucosa using an Interject Injection Needle catheter (Boston Scientific) to separate the mucosal layer from the muscularis. A 20 mm Captivator II snare (Boston Scientific) was then used to excise the mucosal layer, forming a lesion with an exposed submucosa of approximately 20-30 mm in diameter. Piecemeal en bloc resection was performed as needed to achieve the target lesion size. Olympus gastroscope (GIF-2TH180) was used to access and visualize the gastrointestinal tract. For the oozing application, a plain black catheter delivered around 1 ml of material onto the lesion site on the colon right wall under +8 light, followed by curing for 2 min. Adhesion was assessed after 90 min by prodding and shearing the hydrogel with the Interject Needle. Necropsy and H&E staining were performed to evaluate lesion presence, material coverage, and adherence. For the spraying application, the effervescent device was operated at 5 ml/min injection rate and 6 psi CO_2_ pressure. Approximately 1 ml of material was sprayed onto a stomach EMR lesion on the ceiling and cured under +8 light for 2 min. Adhesion was immediately assessed by prodding and pulling the material using the catheter tip and graspers.

***In vivo pig chronic EMR model*:** Four endoscopic mucosal resection (EMR) lesions were created in the colon of one male pig from the proximal to the distal end, using the procedure described above. The first lesion served as a control and was left untreated. For the remaining three lesions, 0.5 – 2 ml of hydrogel precursor solution was delivered onto each site using a plain black catheter under −8 light. In cases where the lesion was difficult to access, the material was applied incrementally using the stamping technique. The light setting was then switched to +8 to cure the material for 2 minutes. Bleeding, material coverage, and adhesion were evaluated in situ via endoscopic visualization.

Animal health was continuously monitored throughout the procedure. On day 3, the pig was euthanized, and necropsy was performed to assess hydrogel retention, mucosal and submucosal adhesion, and to screen for any signs of bowel obstruction, bleeding, perforation, inflammation, or edema. Material adhesion was tested by attempting to peel or dislodge the hydrogel using forceps. Lesion sites were harvested for histological analysis. H&E and Masson’s trichrome staining were performed, and histopathological assessment was conducted to evaluate inflammation, necrosis, fibroplasia, and neovascularization, each graded on a 0–4 scale based on a published protocol (Fig._S19).[126]_

***Statistics*:** All experiments were performed in at least triplicate, and data are presented as mean ± standard deviation (SD). Statistical analyses were conducted using GraphPad Prism (version 10.3.1, www.graphpad.com). Unpaired, two-tailed Student’s *t*-tests were used for comparisons between two groups, and one-way ANOVA with Tukey’s post hoc test was applied for multiple group comparisons. A non-linear variable slope model was used to calculate IC_50_ values from cytotoxicity assays. Statistical significance was defined as P < 0.05 (*P < 0.05, **P < 0.01, ***P < 0.001, ****P < 0.0001; ns = not significant). Some illustrations were created using BioRender.com.

