## supporting information for "In Situ Photoactivated Hydrogel Adhesive Dressings for Post Colon Polypectomies (PolypCures)"

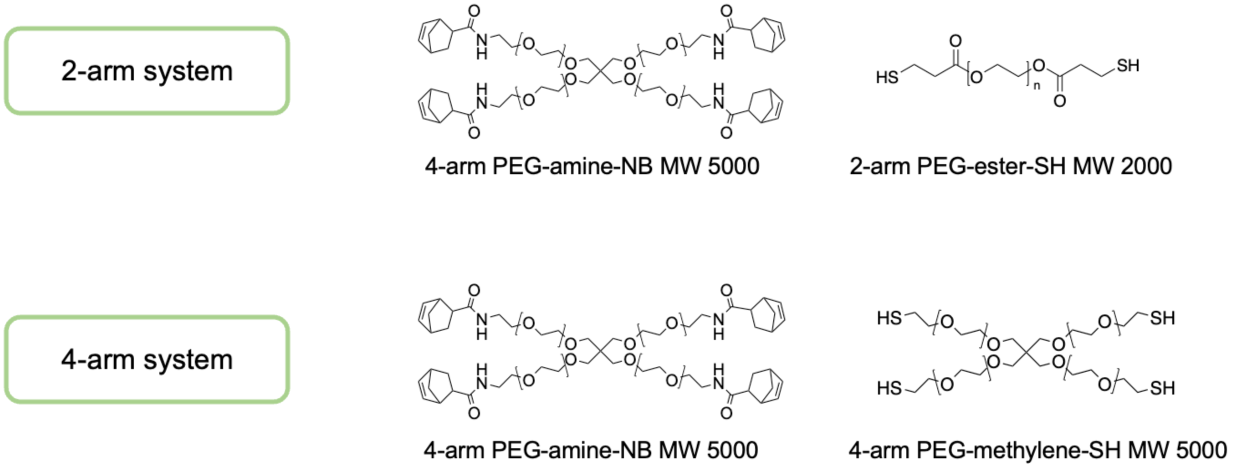


Fig S1. Polymer systems used for hydrogel crosslinking.

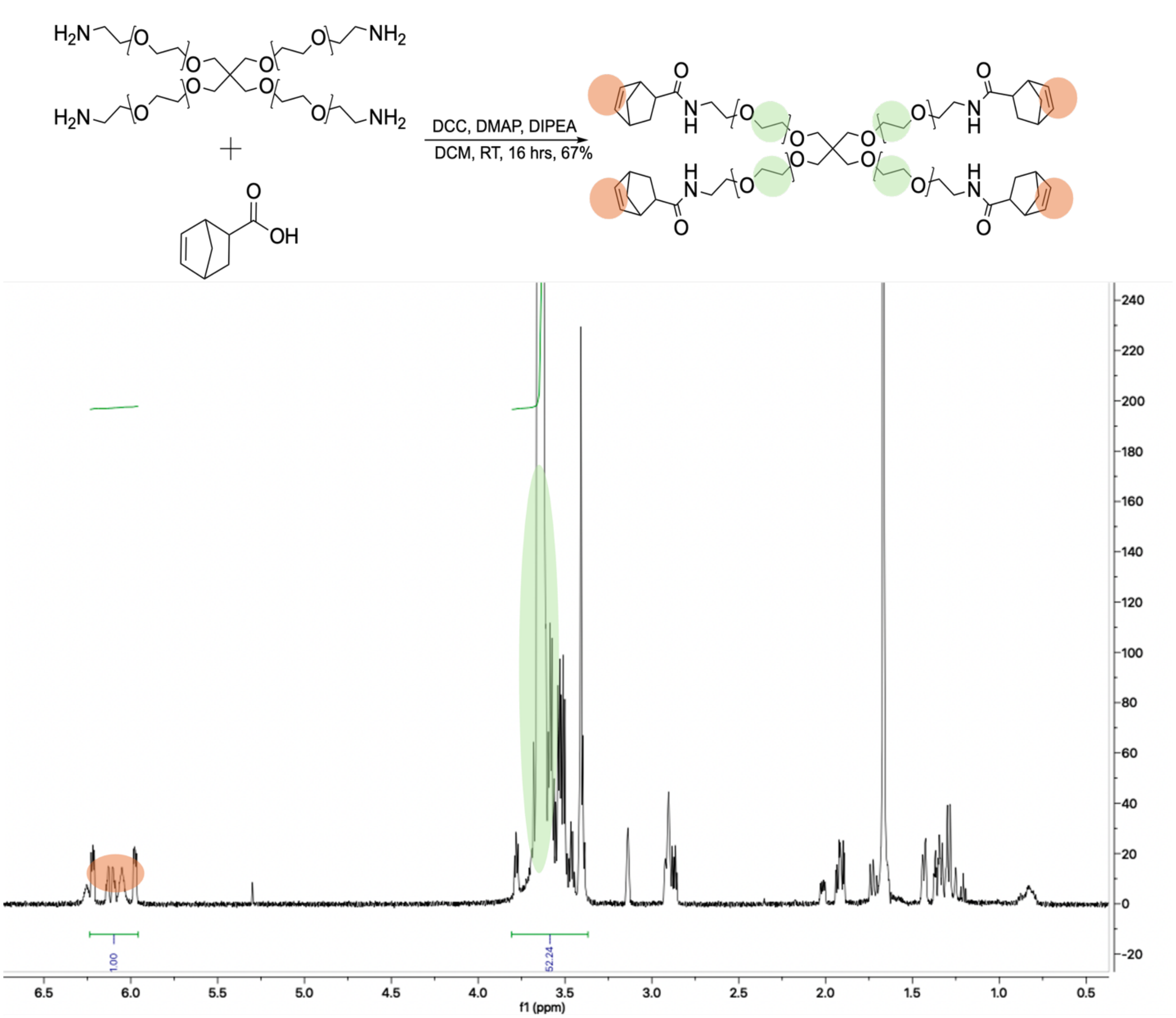


Fig S2. Synthesis scheme and $^{1}$H-NMR of 4-arm PEG-amine-NB MW 5000. This is the functionalization of PEG with NB groups through DCC coupling.

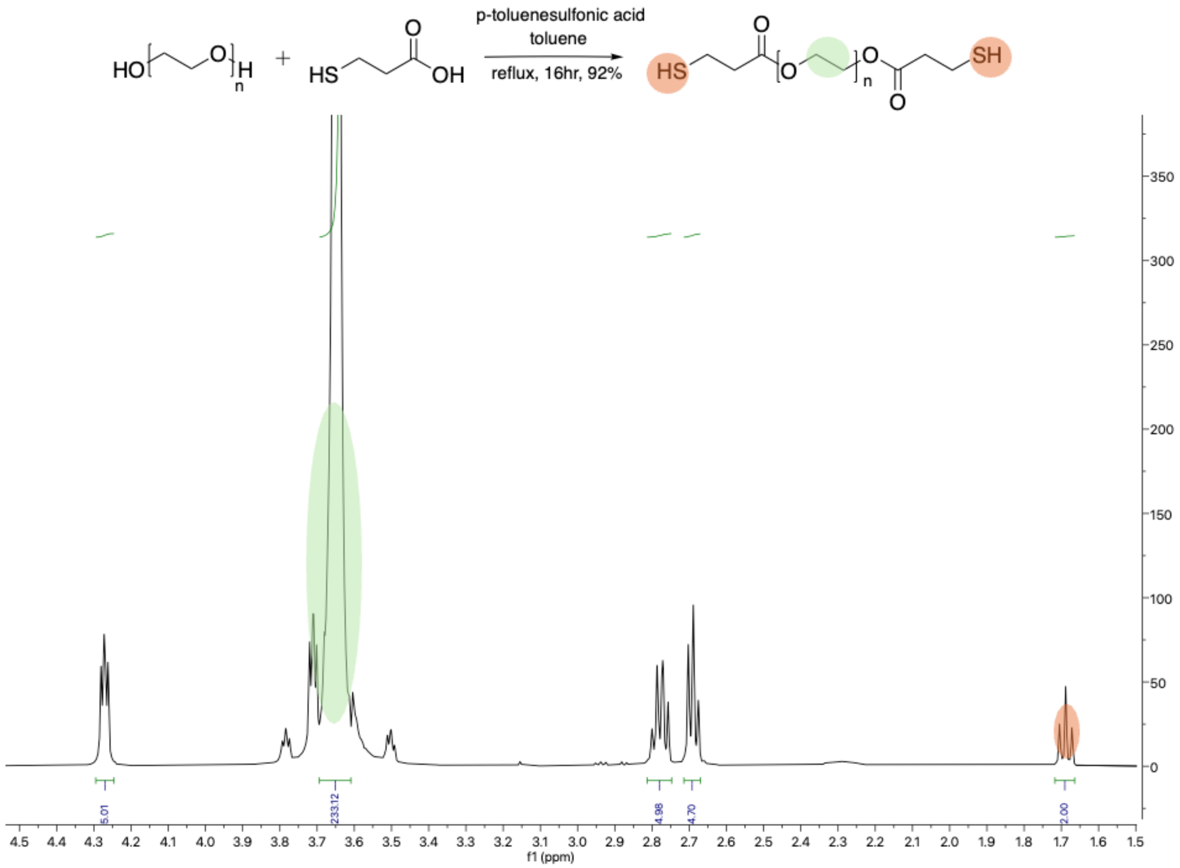


Fig S3. Synthesis scheme and $^{1}$H-NMR of 2-am PEG-ester-SH MW 2000.

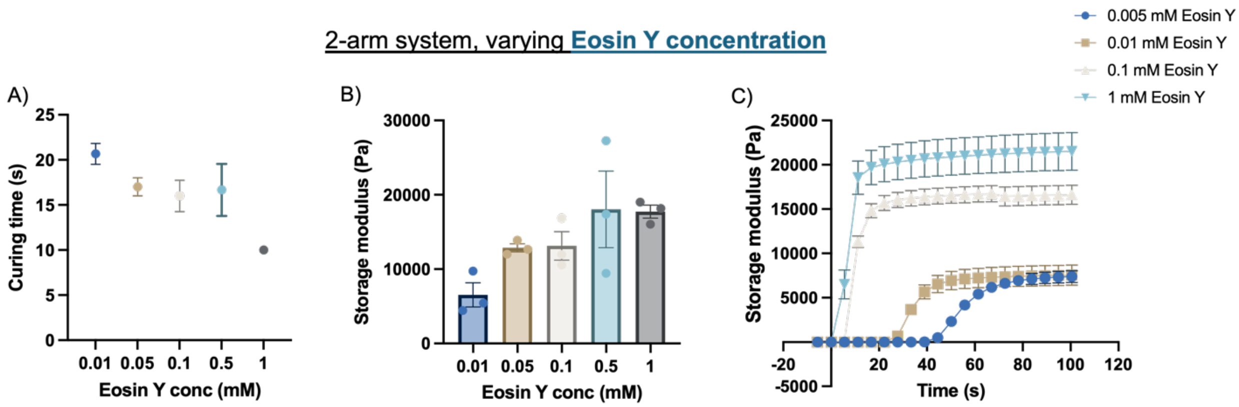


Fig S4. Mechanical characterizations of the 2-arm system. A-C) Curing time and storage modulus measured at varying Eosin Y concentrations for the 150W AmScope Max light initiated photo crosslinking system.

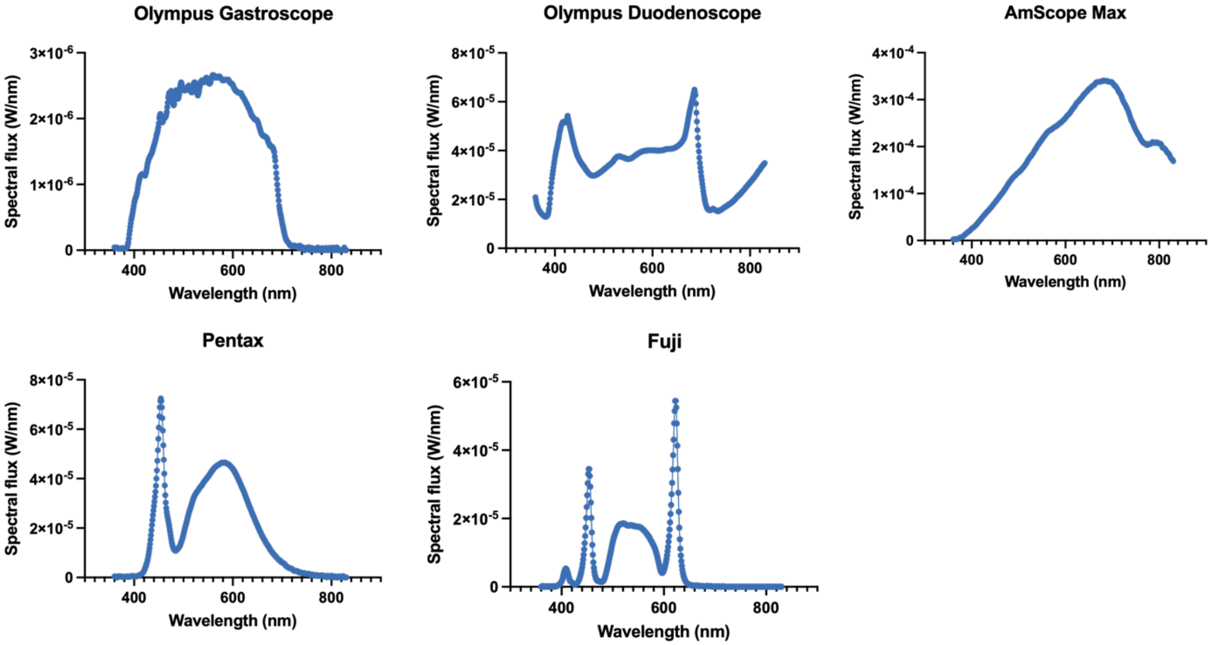


Fig S5. Spectral flux measured at +1 light setting, 2 cm away from light using various light sources.

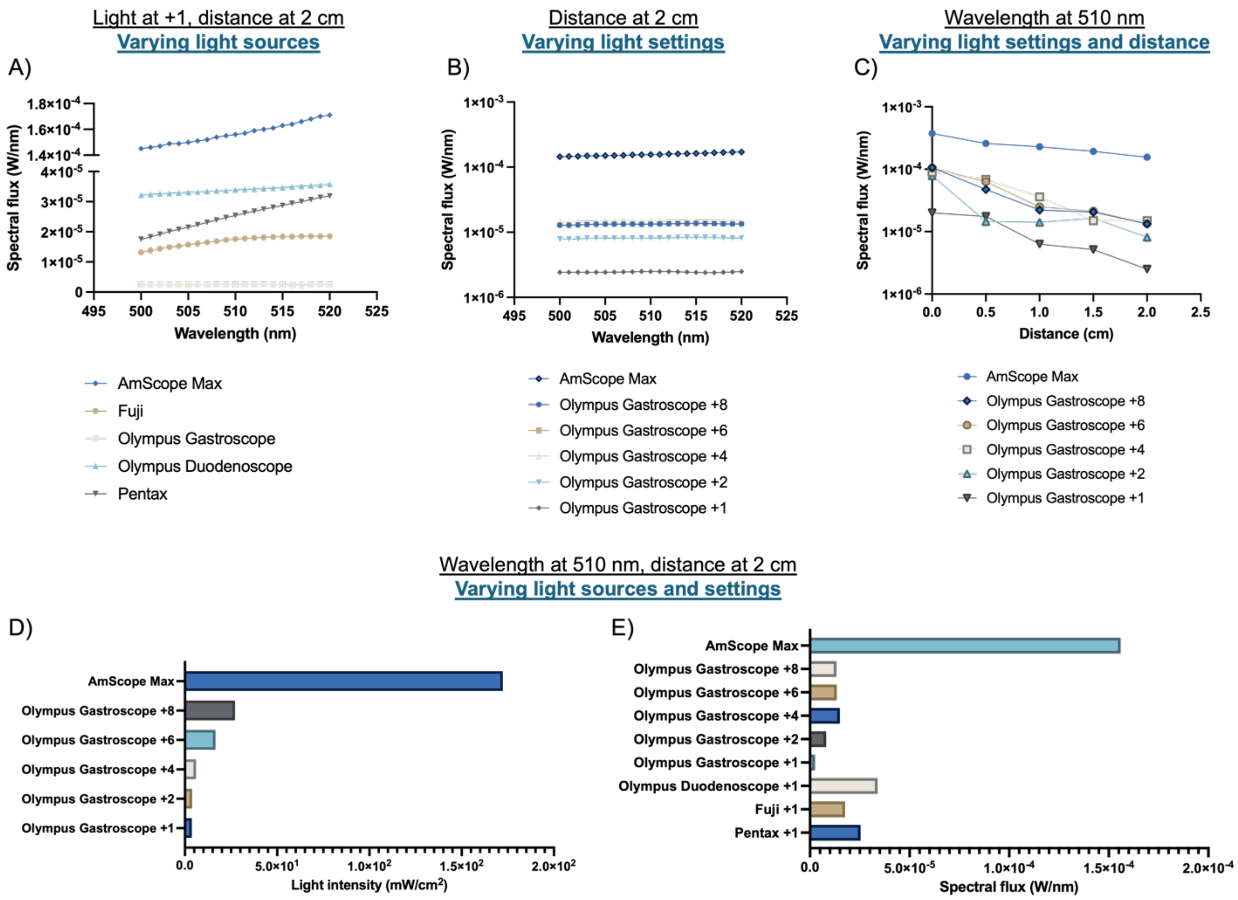


Fig S6. White light characterization of various light sources at wavelengths near 510 nm. 510 nm is the absorbance peak of Eosin Y. A-C) Spectral flux is distance and light setting dependent. D-E) Endoscope light is much weaker than AmScope max light settings in terms of both light intensity and spectral flux.

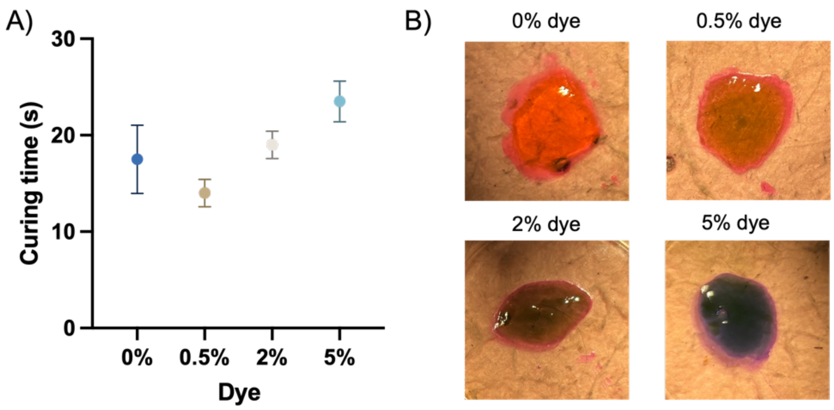


Fig S7. Addition of FD&C blue #1 dye slightly delays curing. 0.5% blue dye is sufficient for visualization and doesn't delay curing. This is the 2-arm system with 1 mM Eosin Y, cured under AmScope max light settings.

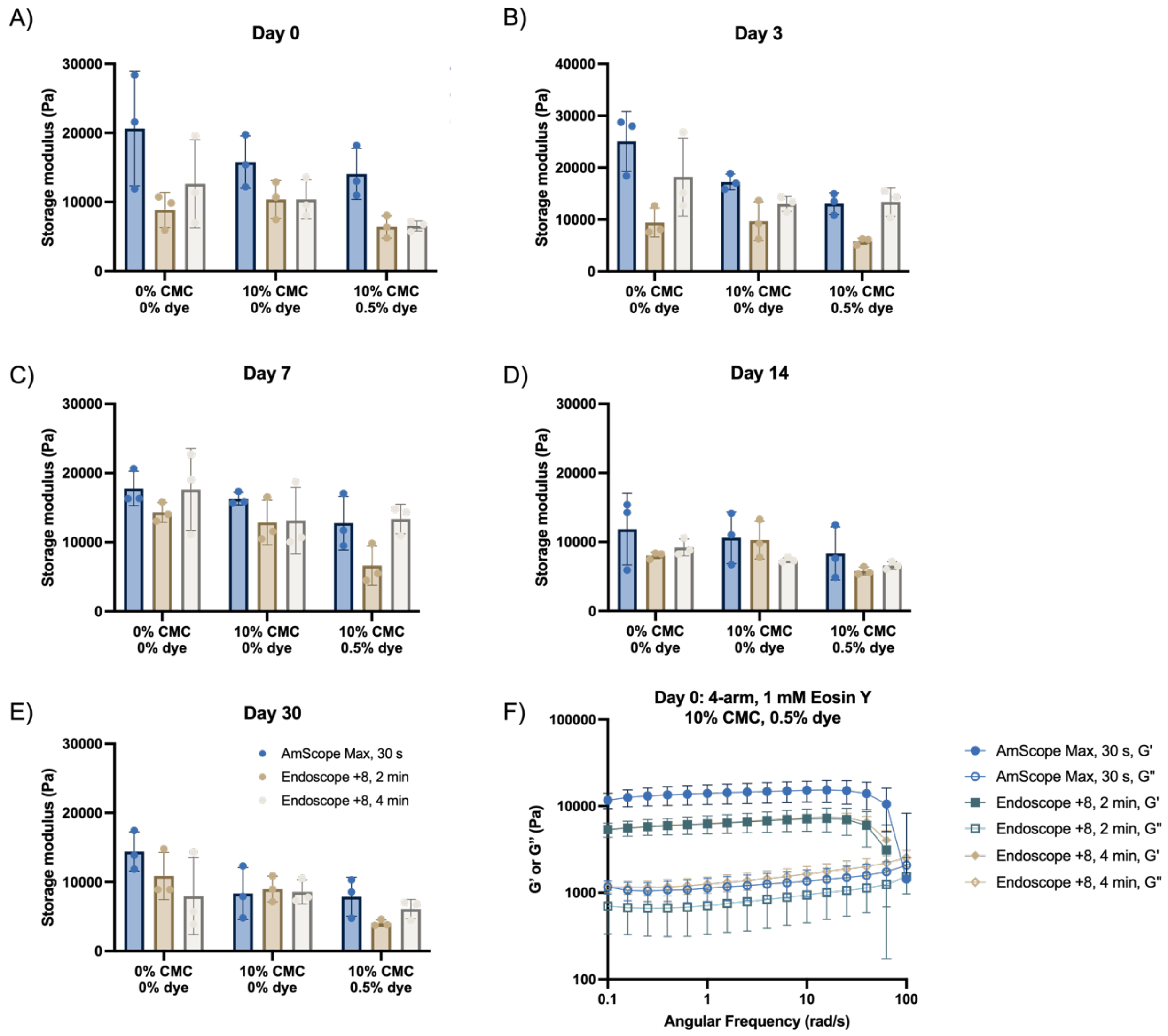


Fig S8. A-E) 4-arm system hydrogels at 1 mM Eosin Y with or without 10% CMC or 0.5% dye cured through either AmScope max light setting or Olympus Gastroscope +8 setting. Hydrogels were submerged in PBS for up to 30 days and measured for their storage modulus over time. F) Storage and loss moduli of the 4-arm system hydrogels at 1 mM Eosin Y with 10% CMC and 0.5% dye cured with various light sources on day 0.

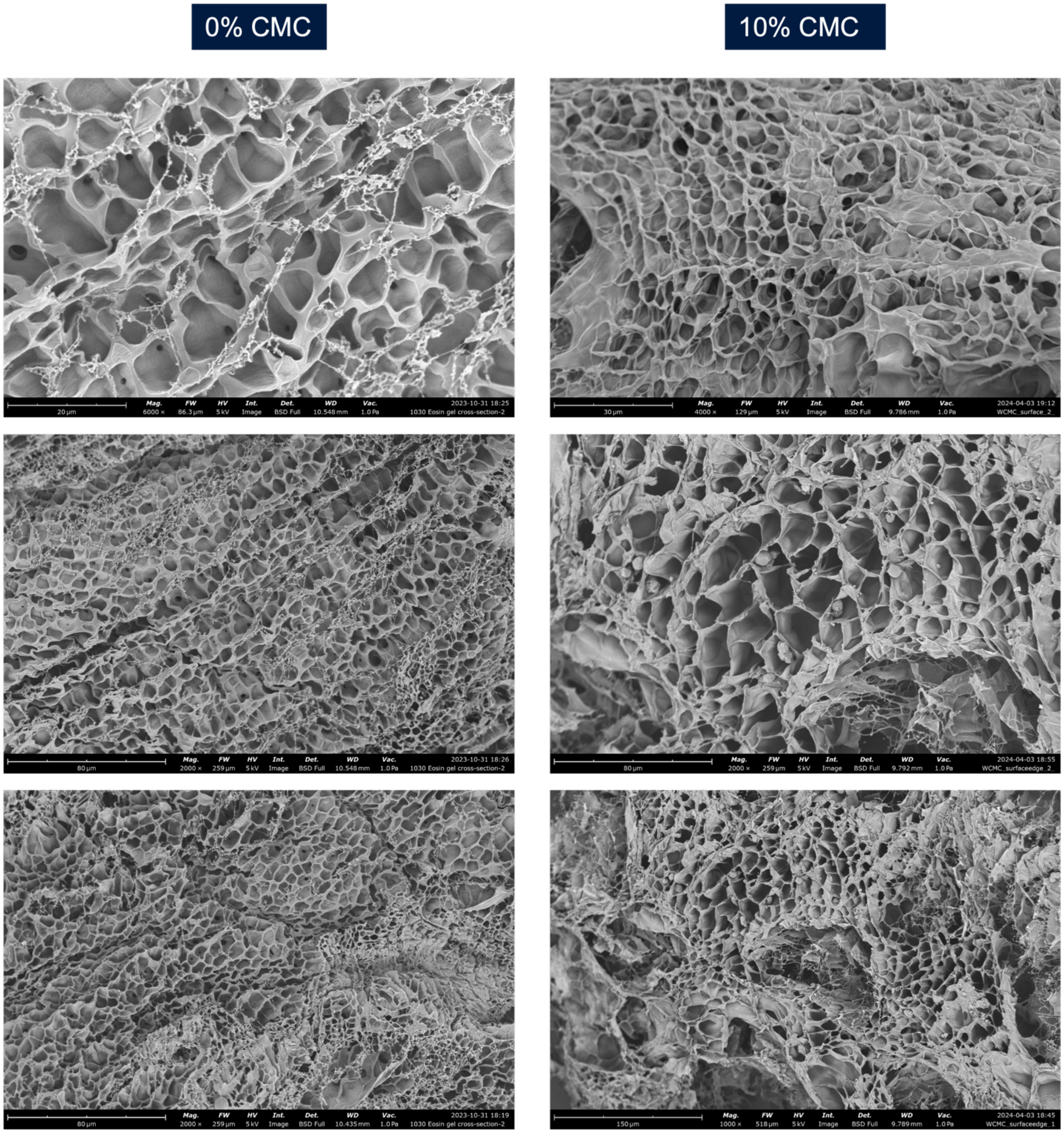


Fig S9. SEM images of the 4-arm system hydrogel at 1 mM Eosin Y cured with Endoscope +8 light, with varying % CMC.

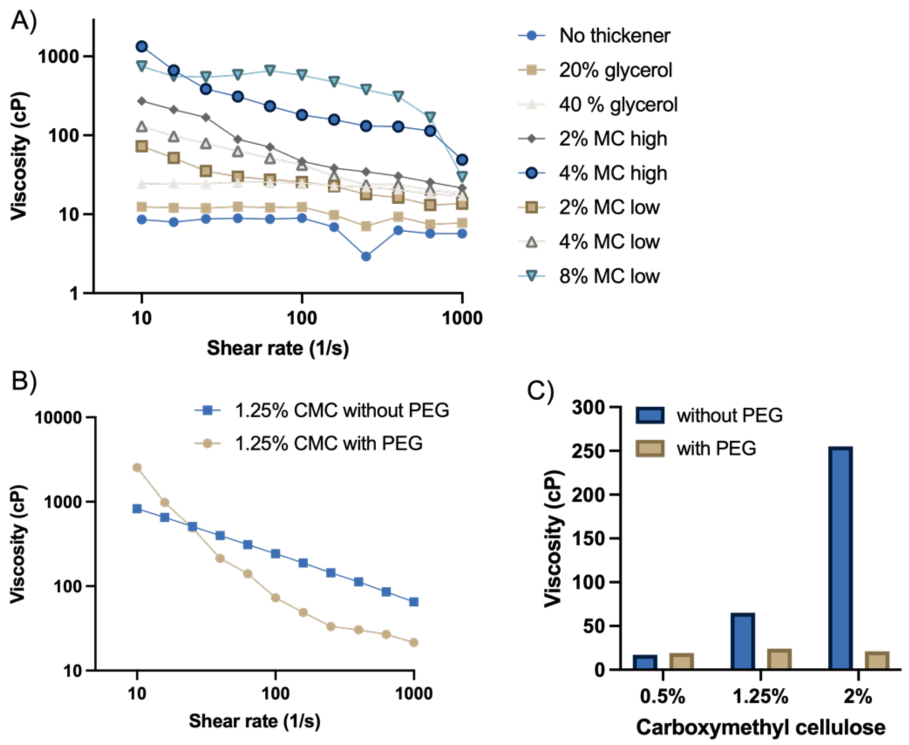


Fig S10. A) Viscosity measurements of varying concentrations of thickeners: glycerol, methyl cellulose 4000 cP (MC high), methyl cellulose 1500 cP (MC low) added to 20 wt% PEG 4000. PEG 4000 act as a substitute for the 4-arm system polymers for larger scale viscosity measurements. High concentrations of MC lead to heterogenous mixing of the solution. B) PEG acts as a plasticizer in the carboxymethyl cellulose (CMC) solution as the shear rate increases. C) Viscosity measured at shear rate $10^3$ $s^{-1}$ for each condition.

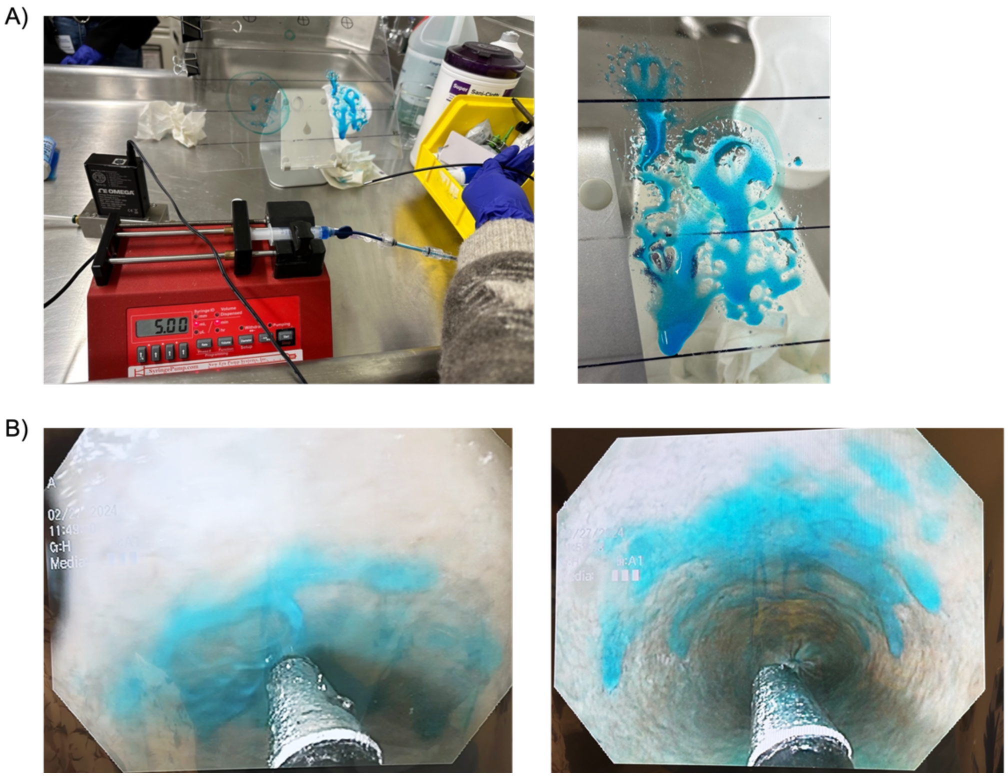


Fig S11. Spraying patterns of the air-assisted device. A) The catheter demonstrated precise targeting capability. B) The device produced a uniform, thin spray layer in the *ex vivo* full colon model. A 1.25% CMC solution with blue dye, chosen for its viscosity similarity to the best-performing hydrogel precursor solution, was used as a dummy material for visualization.

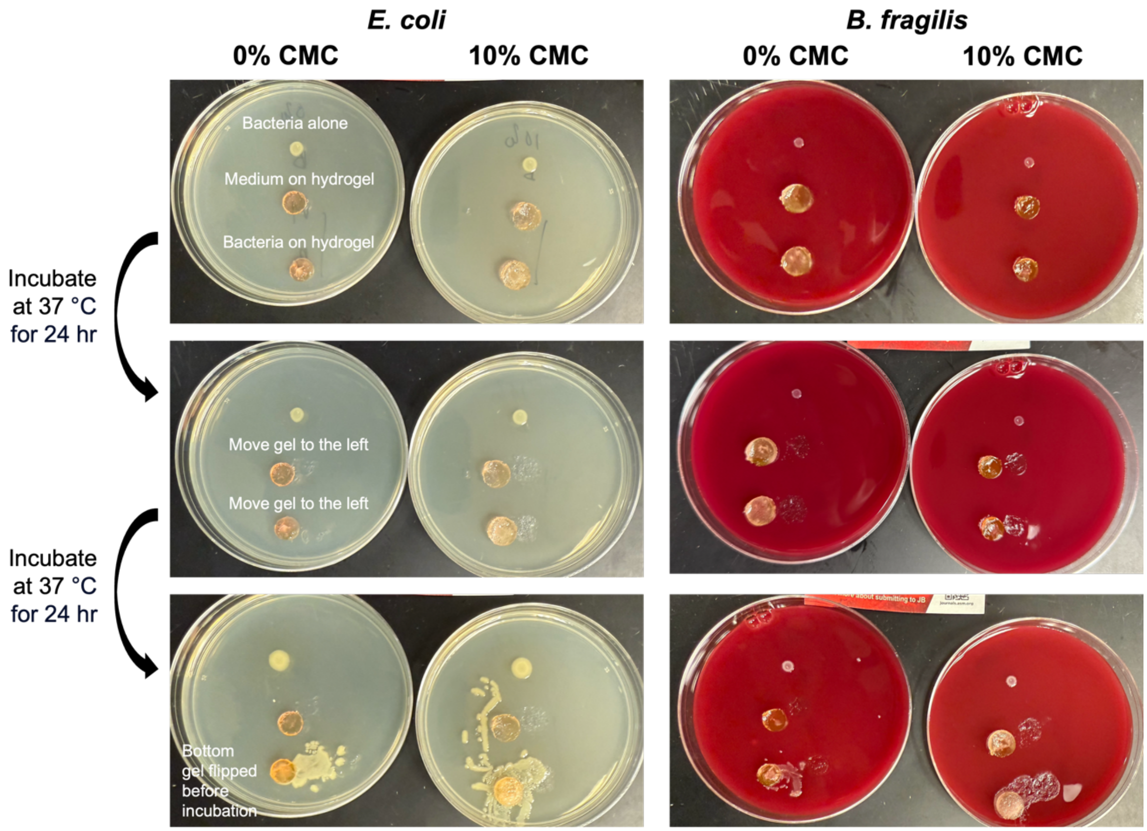


Fig S12. Representative images from the agar plate assay assessing bacterial migration through the hydrogel. Each plate includes three groups: (1) top – bacteria only (positive control), (2) middle – medium on hydrogel (negative control), and (3) bottom – bacteria on hydrogel. After 24 hr incubation at 37 °C, no bacteria growth was observed beneath the hydrogel, indicating effective barrier function against *E. coli* and *B. fragilis* migration through the gel. Hydrogels inoculated with bacteria were then flipped and incubated for an additional 24 hr, confirming bacterial viability on the apical surface.

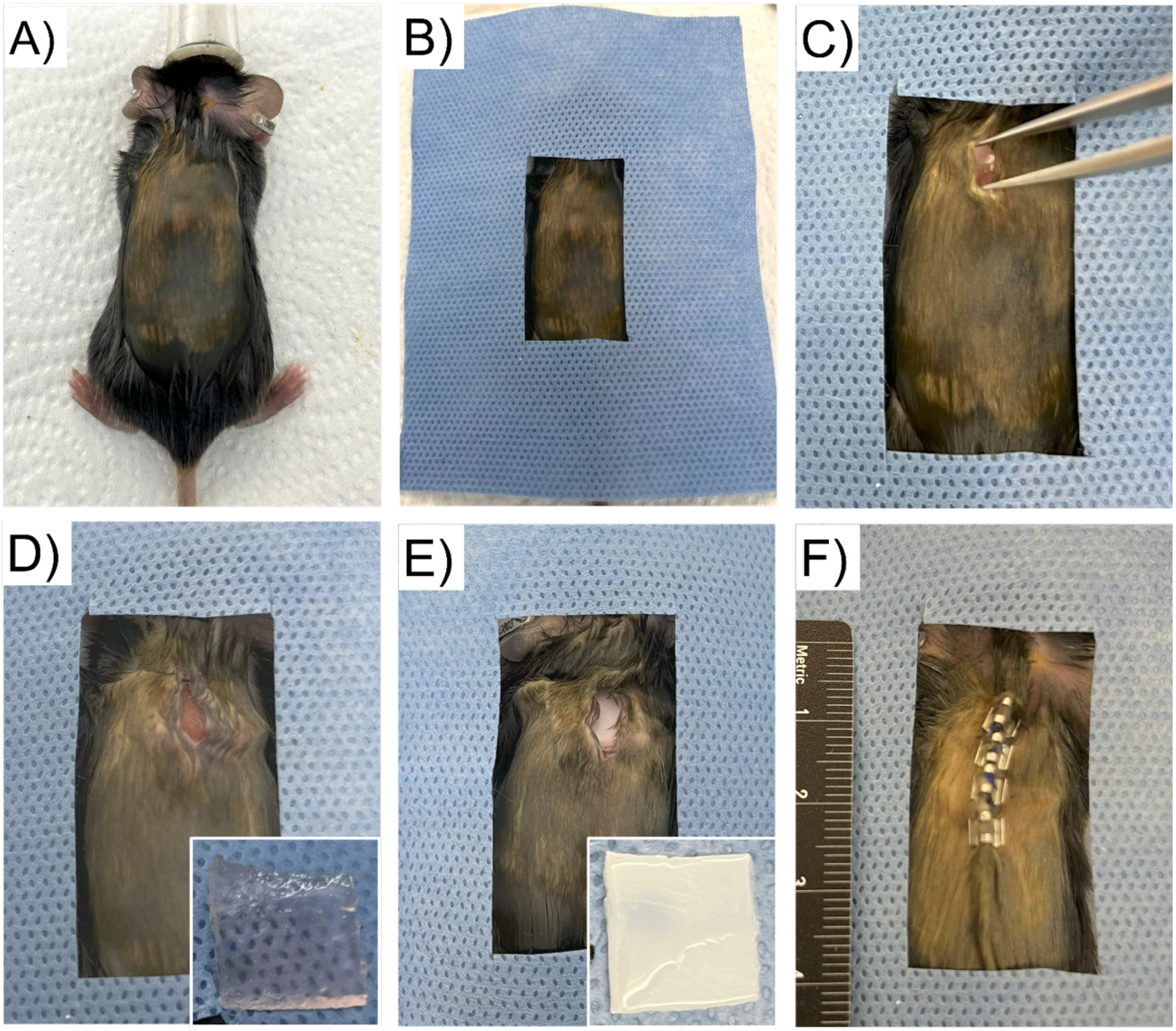


Fig S13. Representative images from the mouse subcutaneous implantation study. A-B) Mice were anesthetized and draped. C) An approximate 1 cm dorsal skin incision was made to create a subcutaneous pocket. D-E) Hydrogels or high-density polyethylene (HDPE) implants (1 x 1 x 0.2 cm) placed in the subcutaneous pocket. Inserts showing implants. F) The incision was closed with four wound clips post-implantation.

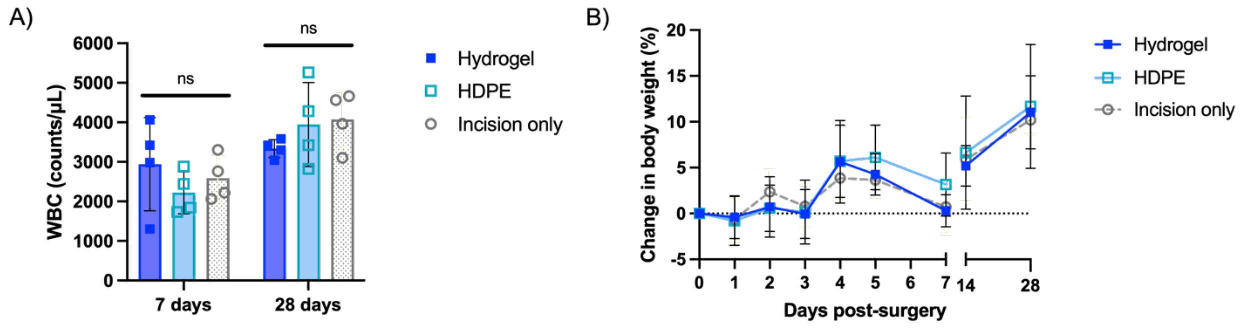


Fig S14. A) White blood cell (WBC) counts were comparable across all groups on day 7 and 28. B) All mice exhibited normal body weight trends throughout the study period.

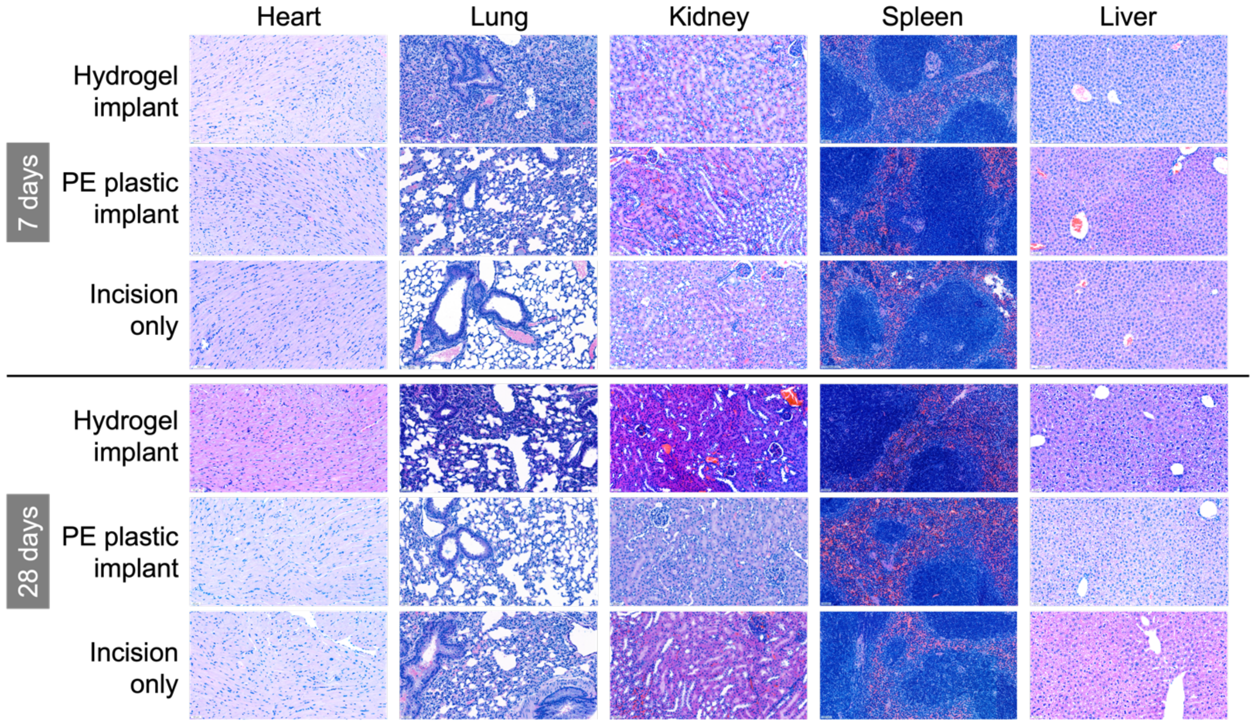


Fig S15. H&E staining of major organs (heart, lung, kidney, spleen, and liver) from the mouse subcutaneous implantation study. No significant pathological abnormalities were observed in any group at either day 7 or day 28.

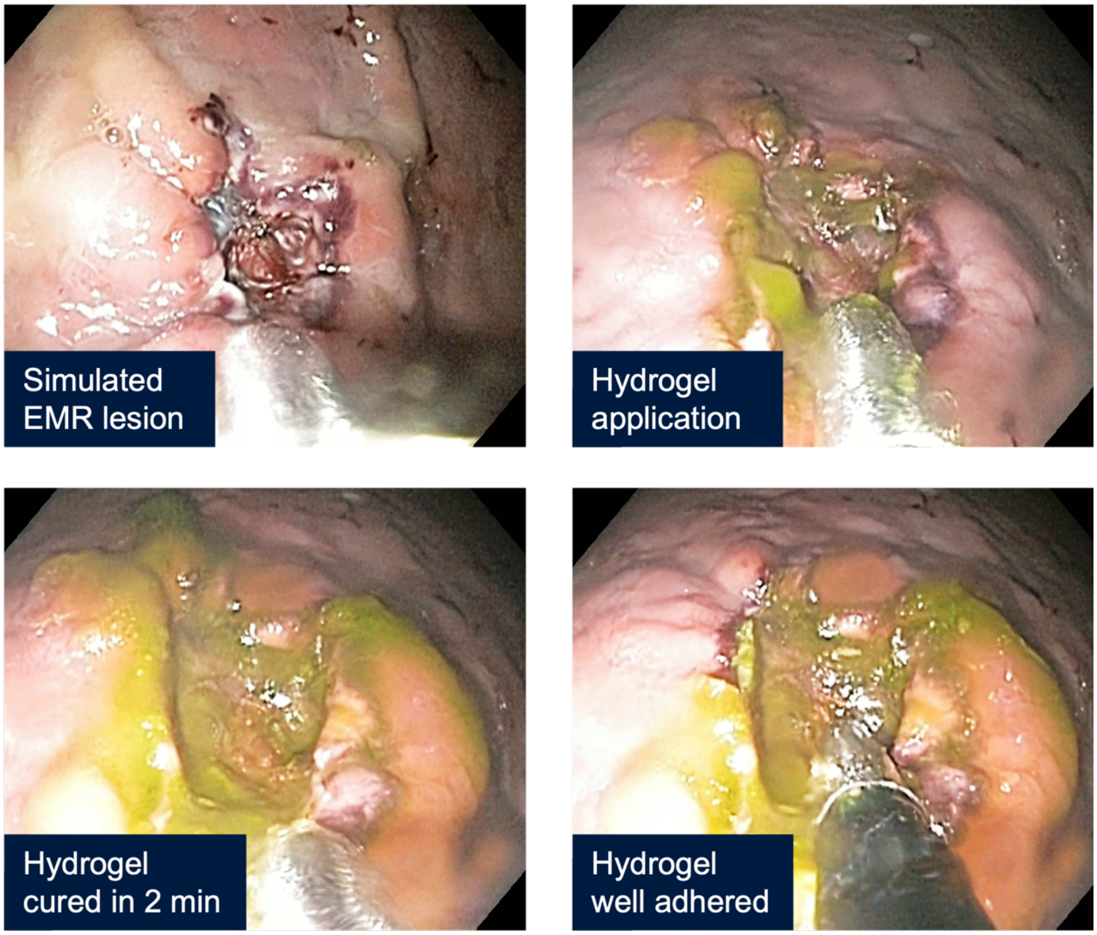


Fig S16. Spraying application of material in an *in vivo* pig stomach acute endoscopic mucosal resection (EMR) model. The material was ejected as a fast atomized stream, forming a thin, uniform layer that fully covered the lesion. After 2 min of curing under +8 light, material adhered strongly to the submucosa. Forceful removal of the material caused bleeding, and even graspers could not fully detach it (Mov S3).

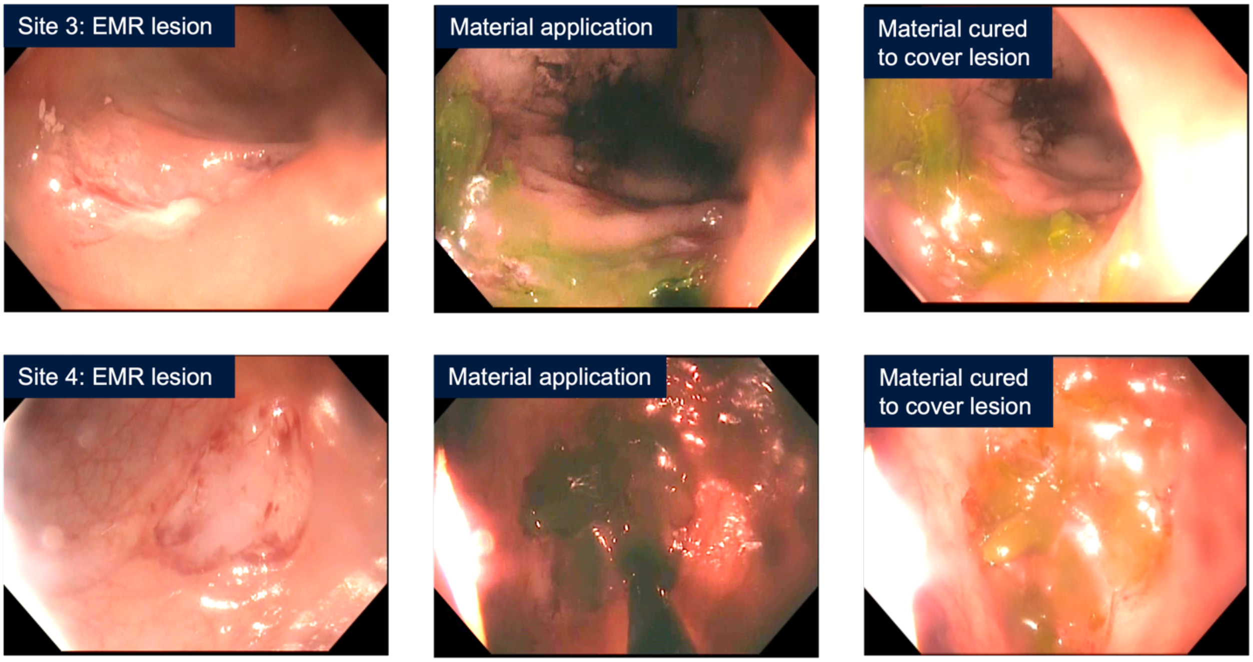


Fig S17. Oozing application of material in a 3-day *in vivo* pig colon chronic endoscopic mucosal resection (EMR) model. Colon lesions measuring 20–30 mm in diameter were created using a cold snare technique. Hydrogel material was applied incrementally via a plain black catheter, oozed directly onto the lesion site under -8 light. The material remained localized at the application site, with no displacement by the catheter during delivery. Following 2 min of curing under +8 light, the hydrogel formed a thin, uniform layer that fully covered the lesion. No bowel obstruction or bleeding was observed post-application (Mov S4).

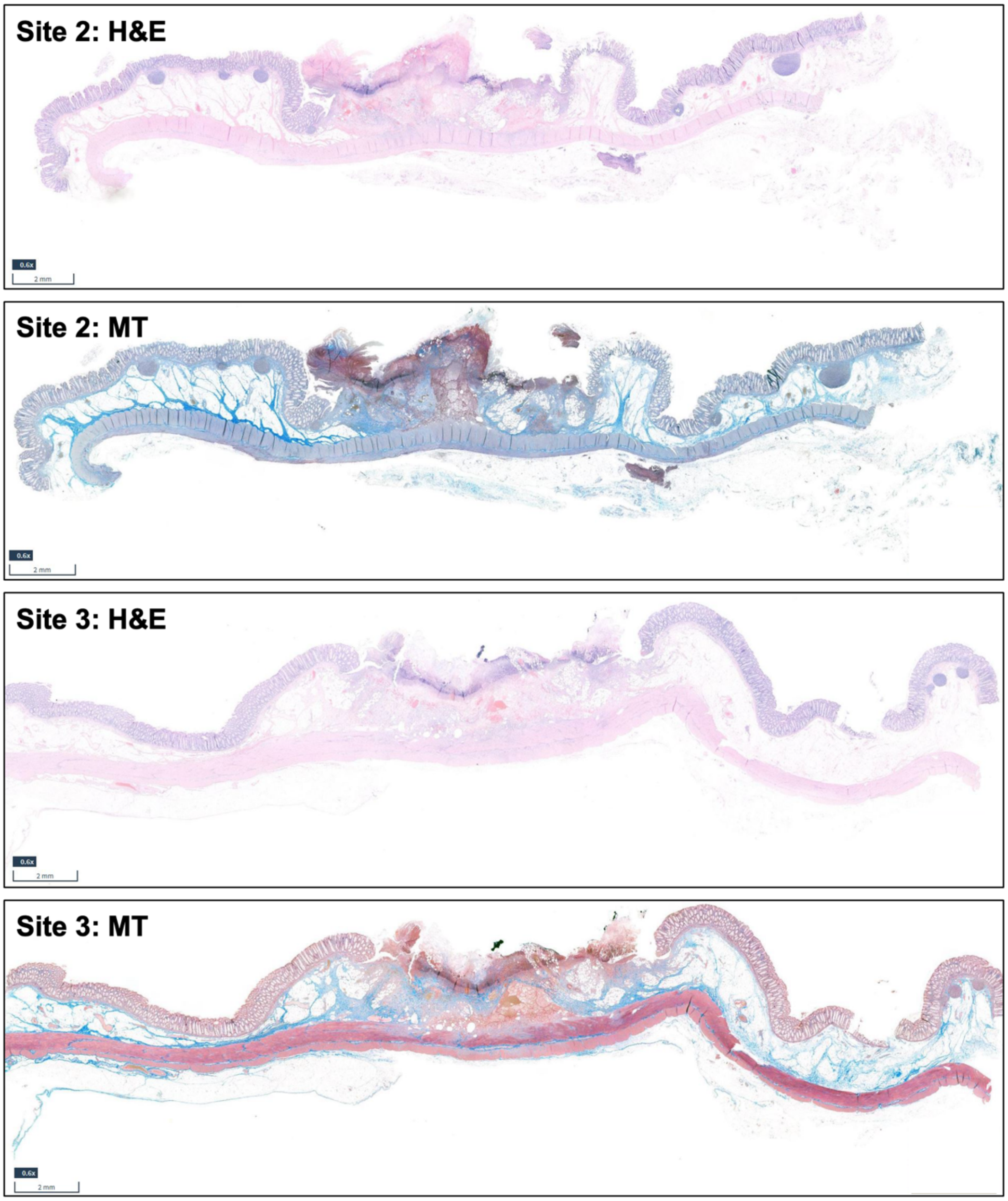


Fig S18. H&E and Masson's trichrome (MT) staining of lesion sites from the 3-day *in vivo* pig chronic EMR model. Site 2 and site 3 were both treated with hydrogel. Staining images of site 1 (control) and site 4 (hydrogel-treated) are shown in Fig. 5D.

| | **Inflammation** | **Necrosis** | **Fibroplasia** | **Neovascularization** |
|---|---|---|---|---|
| **Score 0** | Absent/minimal (<1%) | Absent/minimal (<1%) | Absent/minimal (<1%) | Absent/minimal (<1%) |
| **Score 1** | Mild/occasional inflammatory cells | 1-24% of area showing necrosis | 1-24% of area showing fibroplasia | 1-24% of area showing neovascularization |
| **Score 2** | Moderate (visible increase/small clusters) | 25-49% of area showing necrosis | 25-49% of area showing fibroplasia | 25-49% of area showing neovascularization |
| **Score 3** | Severe (large number of inflammatory cells) | 50-74% of area showing necrosis | 50-74% of area showing fibroplasia | 50-74% of area showing neovascularization |
| **Score 4** | Extensive (with abscess/ulcers) | 75-100% of area showing necrosis | 75-100% of area showing fibroplasia | 75-100% of area showing neovascularization |

Fig S19. Histopathological scoring system used to evaluate tissue responses to hydrogel in the pig chronic EMR model.
